# Structure-function profiling identifies determinants of TCR-T cell therapeutic efficacy

**DOI:** 10.64898/2026.09.15.750028

**Authors:** Leire Oyon-Olea, Enric Vercher, María Gilda Dichiara-Rodríguez, Elena Erausquin, Sandra Hervás, Jacinto López-Sagaseta

## Abstract

Glypican-3 (GPC3) is a promising target in adoptive T-cell therapy for hepatocellular carcinoma (HCC). TCR-A and TCR-B are two recently identified GPC3-specific TCRs recognizing the same HLA-A*02:01-restricted epitope but displaying markedly different therapeutic efficacy. To define the determinants of productive TCR-antigen recognition, we integrated high-resolution structures of both binary pMHC and ternary TCR:pMHC complexes, biolayer interferometry, peptide mutagenesis, target-cell conjugation assays, repetitive antigen challenge (RAC) and in vivo models. Structurally, conformational permissiveness of the immunodominant GPC3(522-530) peptide within HLA-A*02:01’s groove is central for productive TCR docking, with TCR-B presenting slower association but prolonged dwell time. Functional divergence emerged during target-cell engagement and amplified under RAC conditions, where TCR-B sustained cytotoxic activity while TCR-A progressively lost function. This superior functional endurance translated into complete tumor eradication and durable responses in vivo. These findings provide a mechanistic basis and a conceptual framework for the selection and optimization of TCRs for adoptive immunotherapy.

## Introduction

Hepatocellular carcinoma (HCC) remains a major therapeutic challenge, particularly for patients with advanced-stage disease or those showing limited response to current systemic treatments^1,2^.

While immunotherapy has provided significant advances in specific subsets of patients with HCC^2–4^, current therapeutic strategies remain insufficiently effective, and further approaches are needed. One such approach is adoptive T-cell therapy, a strategy that redirects cytotoxic T cells to target tumor-associated antigens^5^. In solid tumors, however, its therapeutic efficacy depends not only on the molecular specificity driving antigen recognition by the T-cell receptor (TCR). Persistence of engineered T cells and the capacity to successively engage tumor cells and preserve effector function under sustained antigen exposure are also key properties that ultimately determine the efficacy of the adoptive T-cell therapy^5,6^.

Glypican-3 (GPC3), an oncofetal cell-membrane glycoprotein is abundantly expressed in HCC and other embryonal or epithelial tumors^7,8^. Importantly, GPC3 shows reduced expression in most healthy and adult tissues. Altogether, GPC3 represents a highly relevant antigen for HCC immunotherapy^7–9^.

Earlier GPC3-directed CAR-T approaches showed limited clinical efficacy, with insufficient persistence, T-cell dysfunction and loss or shedding of surface GPC3^9,10^, although newer engineered CAR-T platforms have produced encouraging early clinical activity^11^. An alternative approach is provided by T cells engineered with antigen-specific TCRs^12,13^. Unlike CARs, TCRs recognize peptides originated intracellularly and presented by antigen presenting molecules^13^. This mechanism provides TCRs the advantage of binding peptides derived from either intracellular, surface and secreted antigens, thus extending the target spectrum beyond cell surface markers^13^.

A 9 amino acid-long motif, with sequence FLAELAYDL and encompassing GPC3 residues 522-530 was originally identified as an HLA-A*02:01-restricted and naturally processed cytotoxic T-lymphocyte peptide antigen^14^. In a recent study, Vercher and colleagues challenged mice with human GPC3 and isolated single clones of murine T lymphocytes that specifically recognized the GPC3(522-530) 9-mer peptide in an HLA-A*02:01-restricted manner^12^. The authors showed that when expressed in primary human T lymphocytes, some of these TCRs drove T lymphocytes towards GPC3^+^ HCC cells. One TCR, TCR-B, displayed superior antitumor properties: enhanced effector and proliferative capacity, persistence, greater tumor infiltration and overall efficacy in xenograft mouse models, outperforming a second-generation 4-1BB-based GPC3 CAR-T approach^12^. Further, NSG (NOD scid gamma) mice bearing HCC tumors (HepG2 or PLC/PRF/5-A2) treated with engineered primary human T cells displaying TCR-B showed tumor clearance and prolonged survival in an IL-2-dependent manner. However, in addition to TCR-B, the same study identified additional GPC3-9-mer-specific TCR clones, such as TCR-A, that displayed comparatively limited antitumoral activity ^12^.

These findings led to a central mechanistic question: why do two TCRs that recognize the same GPC3 immunodominant peptide produce diverse therapeutic profiles?

Here, we compare two GPC3-9-mer-specific TCRs - TCR-A and TCR-B - with different therapeutic efficacies. Both receptors share an identical TCRβ chain and differ exclusively in the CDR3 region of the α chain. While both TCRs target the same GPC3 motif, TCR-A promotes early cytotoxicity, whereas TCR-B mediates more persistent tumor control upon serial antigen challenge. Binding studies, however, did not correlate with a markedly stronger affinity as the basis for TCR-B superiority at the functional level; TCR-B showed superior therapeutic efficacy despite the very low affinity, similar to that of TCR-A, exhibited for HLA-A2:GPC3-9-mer in biolayer interferometry measurements. We also determined high-resolution molecular structures of HLA-A2:GPC3-9-mer and HLA-A2:GPC3-11-mer, as well as the corresponding ternary TCR-A:HLA-A2:GPC3-9-mer and TCR-B:HLA-A2:GPC3-9-mer complexes. Importantly, these structural studies reveal conformational permissiveness of GPC3-9-mer as a key determinant for TCR docking. TCR-A and TCR-B present highly conserved docking footprints, yet subtle local differences in pMHC engagement may contribute to the divergent cellular function.

Collectively, these findings support the implementation of metrics according to cellular persistence, performance under serial antigen challenge and target-cell conjugate steadiness, in combination with structural and biophysical approaches, for selection of optimal TCRs in solid tumor adoptive T-cell therapy.

## Results

### Alternative conformations of HLA-A2-bound GPC3**-**9- and 11-mer peptides confer different pMHC topologies

High-resolution crystal structures were obtained for HLA-A2 bound to 9-mer and 11-mer peptides (Figure 1, Supplementary Figure 1,2 and Supplementary Table 1). The 9-mer peptide adopts a classical HLA-A2-binding mode, with anchor residues located across the peptide-binding groove and a central region with solvent-exposed residues accessible for TCR engagement (Fig. 1a, Supplementary Figure 1 and Supplementary Table 2). The HLA-A2-bound 11-mer stands out for a large and central protruding fold, while preserving the anchoring sites within the HLA-A2 groove (Fig. 1a, Supplementary Figure 2 and Supplementary Table 3). Structural superposition of both complexes shows that the 11-mer conserves the N-terminal core observed for the 9-mer but the additional C-terminal residues lead to a shifted peptide register and pMHC topology (Fig. 1c-e). Regardless of the conformational deviation observed for the 11-mer peptide, it is found bound to the MHC’s groove, structurally confirming HLA-A2’s competence to present both peptide formats. Several intermolecular contacts with the 9-mer peptide are conserved in the HLA-A2:GPC3-11-mer complex, whereas other interactions are lost or reconfigured (Fig. 1d,e and Supplementary Tables 2,3). The 11-mer produces novel polar contacts that involve residues near the peptide C terminus, including interactions with the MHC platform around the peptide-binding groove, such as those established by R65, D77, Y84, T143, and W147 (Fig. 1f and Supplementary Table 3). These results demonstrate that the 9- and 11-mer peptides lead to different pMHC topologies that likely determine TCR recognition.

**Fig. 1:**
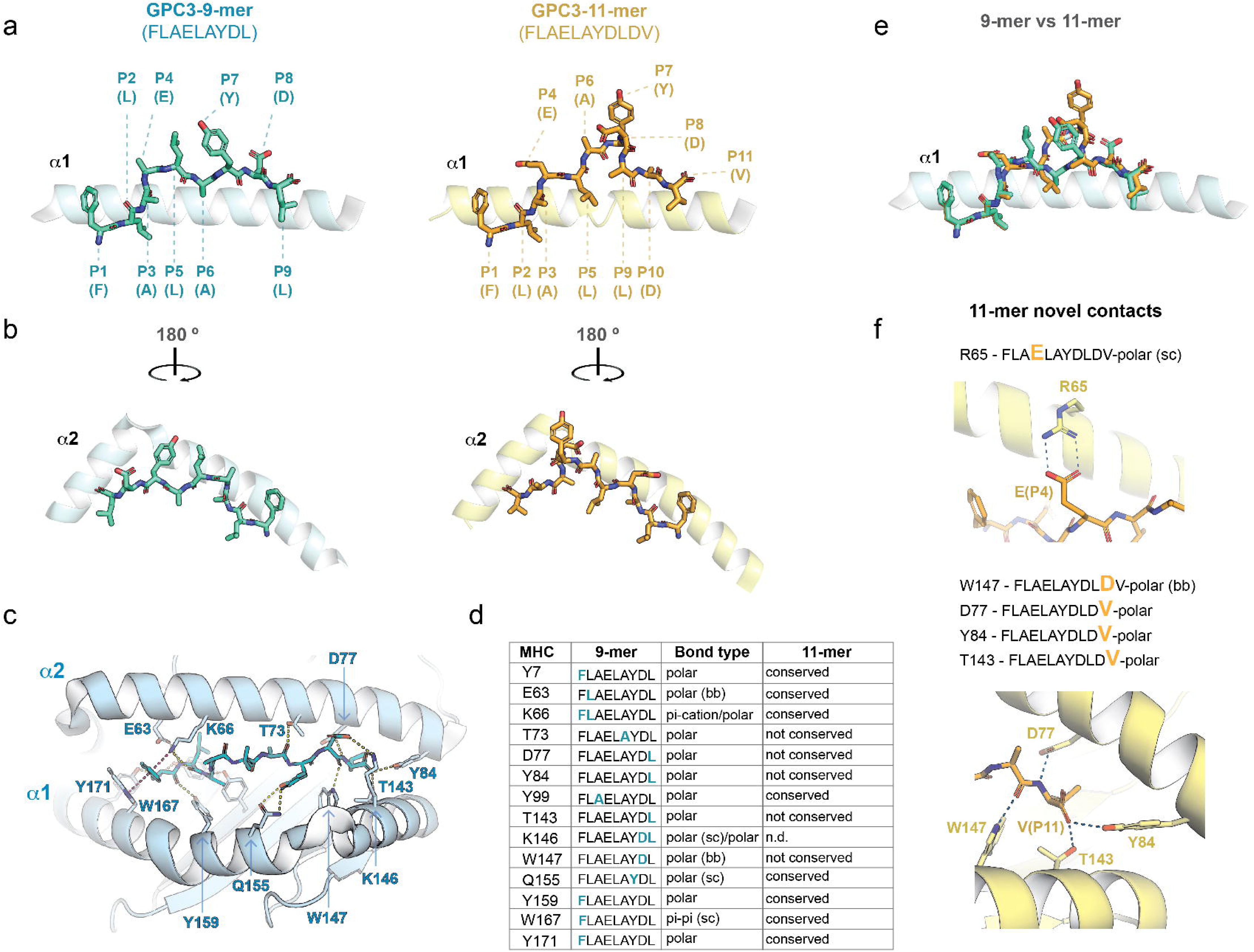
The GPC3 9- and 11-mer peptides adopt different conformations within the HLA-A2 groove. a, Side views of HLA-A2 crystal structures bound to GPC3-9-mer peptide (FLAELAYDL) or the longer 11-mer peptide (FLAELAYDLDV), relative to the HLA-A2 α1 helix. Peptides are displayed as sticks, with the 9-mer in cyan and the 11-mer in orange, inside the HLA-A2 groove, shown in cartoon mode. The position and single-letter code are indicated for each peptide residue. **b,** Alternative views of the HLA-A2:peptide complexes shown in **a** upon 180° rotation around the y axis. **c,** Interaction network mapped for the HLA-A2:GPC3-9-mer complex. Intermolecular contacts are labeled with purple or yellow dashed lines for pi and polar interactions, respectively. **d**, List of HLA-A2:peptide pi and polar interactions found with the 9-mer peptide. Conserved and non-conserved contacts observed with the 11-mer peptide are indicated in the last column. **e,** Structural overlay, relative to the MHC molecule, of the 9- and 11-mer peptides in their HLA-A2-bound states. **f,** Unique polar contacts observed for the 11-mer peptide within HLA-A2. *sc, side chain; bb, backbone; n.d., not determined*.

### Conformational permissiveness of the 9-mer peptide enables TCR engagement through highly analogous docking modes and binding footprints

We next investigated the binding signatures that support GPC3-9-mer recognition by TCR-A and TCR-B. To this end, we performed additional structural studies, crystallized the TCR-bound pMHC complexes, and determined their structures (Supplementary Table 1).

When aligned structurally on the MHC, we first observed a significant rotation of GPC3-9-mer Leu5 (FLAE**<u>L</u>**AYDL) (Fig. 2a and Supplementary Figure 1) in the ternary TCR-A/B:HLA-A2:GPC3-9-mer complexes, relative to the TCR-unbound HLA-A2:GPC3-9-mer structure. In the TCR-unbound state, Leu5 side chain is aligned between the MHC α1 and α2 helices and pointing upwardly, whereas in the TCR-bound form Leu5 appears repositioned through a pronounced backbone rotation. This alternative conformation brings Leu5 side chain against the MHC α2 helix, a shift that translates into a gateway for CDR3β, and therefore, TCR docking onto the pMHC (Fig. 2a). Whether it represents a pre-existing conformation or a receptor-induced alternative fold, this observation reveals that the GPC3-9-mer can adopt different configurations within the HLA-A2 groove, only one of which enables a coherent TCR binding.

**Fig. 2:**
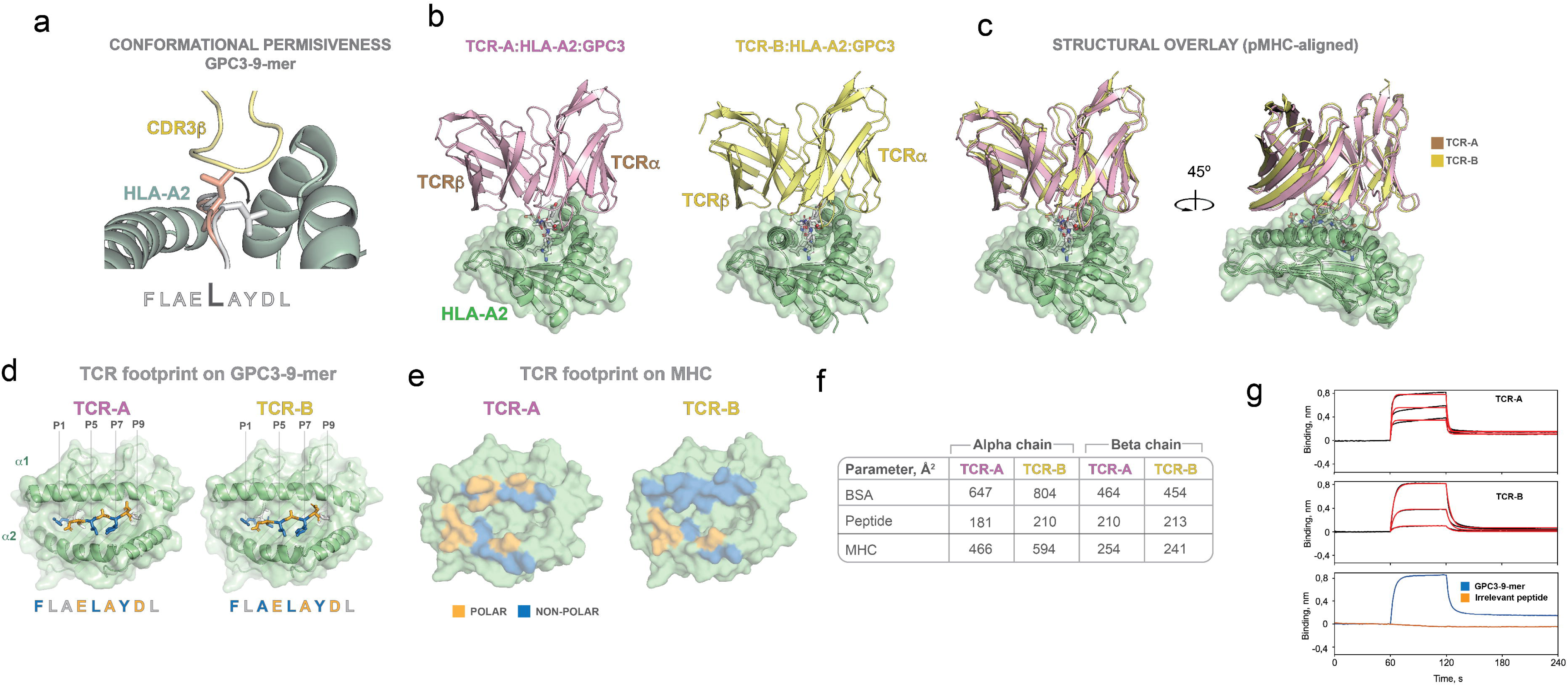
TCR docking is enabled by antigen conformational permissiveness and follows similar pMHC footprints but distinct kinetic profiles. a, Close-up snapshots of GPC3-9-mer peptide as is found bound to only HLA-A2 (pink color) or to TCR-B in the ternary complex (light grey). Conformational permissiveness is highlighted with sticks for Leu5 and a black arrow indicating the rotation observed for this residue upon TCR binding. **b,** The structures of TCR-A (light pink) and TCR-B (yellow), respectively, bound to HLA-A2:GPC3-9-mer are displayed with the MHC molecule shown in green cartoon and surface mode. The GPC3-9-mer antigen is depicted as sticks. For clarity, the TCRs constant regions, the MHC α3 domain and β2 microglobulin are omitted. **c,** Two alternative views of a structural overlay of both TCR:pMHC crystal structures. **d,** The TCR footprints on both the peptide and **e,** the MHC platform. Residues establishing contacts are labeled in either orange (polar contacts) or blue (non-polar contacts) color, and highlighted with sticks in the case of the peptide. The binding footprint on the MHC is shown on a surface representation of the HLA-A2 molecule. **f,** The contribution to the total TCR:pMHC buried surface area is displayed for each TCR chain, and distributed according to either TCR:peptide or TCR:MHC contacts. **g,** Real-time binding assays showing the association and dissociation of TCR-A and TCR-B to varied HLA-A2:peptide complexes. The fitting to a 1:1 Langmuir binding model is indicated with red color. The concentrations of soluble pMHC used in the binding kinetics were 15, 30 and 60 micromolar.

Both TCRs target the HLA-A2:GPC3-9-mer complex in a highly similar and canonical docking manner, sitting on top of the pMHC surface (Fig. 2b and Supplementary Tables 4-13). Structural overlay of the TCR-A:HLA-A2:GPC3 and TCR-B:HLA-A2:GPC3 complexes reveals a notable conservation of the overall docking mode, with both TCRs bound around the central region shaped by the peptide-binding groove (Fig. 2c).

Regardless of the conserved docking interfaces, the footprints on the peptide is largely conserved between the TCRs, whereas the footprint on the MHC platform highlights some differences (Fig. 2d,e). TCR-A and TCR-B form polar contacts with the side chains of acidic residues of the 9-mer peptide located in P4 and P8 (FLA<u>E</u>LAY<u>D</u>L), and with the Ala backbone found in P6 (Fig. 2d). By contrast, the footprint on the MHC platform includes residues found on both α1 and α2 helices that establish both polar and non-polar contacts, but with a distinct distribution between the two TCRs (Fig. 2e). This asymmetry is also captured by the buried surface area analysis. The degree of involvement of the TCRs alpha and beta chains is not equal in all instances. In particular, the TCR-B alpha chain contributes in a large proportion, relative to TCR-A, to the overall BSA, while the beta chain contribution remains comparable between both receptors (Fig. 2f).

We performed biolayer interferometry binding assays to monitor the kinetics that underlie recognition of HLA-A2:GPC3-9-mer by TCR-A and TCR-B. TCR-A presented a fast association and dissociation kinetic profile, whereas TCR-B showed slower on- and off-rates (Fig. 2g and Supplementary Table 14). This suggests that docking of TCR-B on HLA-A2:GPC3-9-mer is slower, but once bound, the complex is more stable than that formed by its TCR-A counterpart. In both cases, the overall affinities fall in the low micromolar range (K_D_ 300-400 μM), thus representing rather very low affinity recognition profiles. The absence of detectable association to HLA-A2 refolded with an irrelevant peptide confirmed the molecular specificity associated to recognition of HLA-A2:GPC3-9-mer (Fig. 2g).

Together, these structural data identify conformational permissiveness of the GPC3 peptide as a prerequisite for productive TCR engagement, while indicating that both receptors recognize a similar pMHC architecture once this permissive state is achieved, accompanied by very low-affinity binding modes.

### GPC3 recognition is driven by conserved and TCR-specific contacts

A close inspection of the TCR:pMHC binding interface revealed the intermolecular interactions that govern the recognition of the MHC-bound 9-mer peptide (Supplementary Tables 4-13). As noted above, amongst the conserved contacts, polar interactions are formed with residues E4, A6 and D8 in the 9-mer peptide, indicating that both TCR footprints share a specific antigenic motif that concentrates the core structural determinant for GPC3-9-mer recognition (Fig. 3a, Supplementary Tables 4-13).

**Fig. 3:**
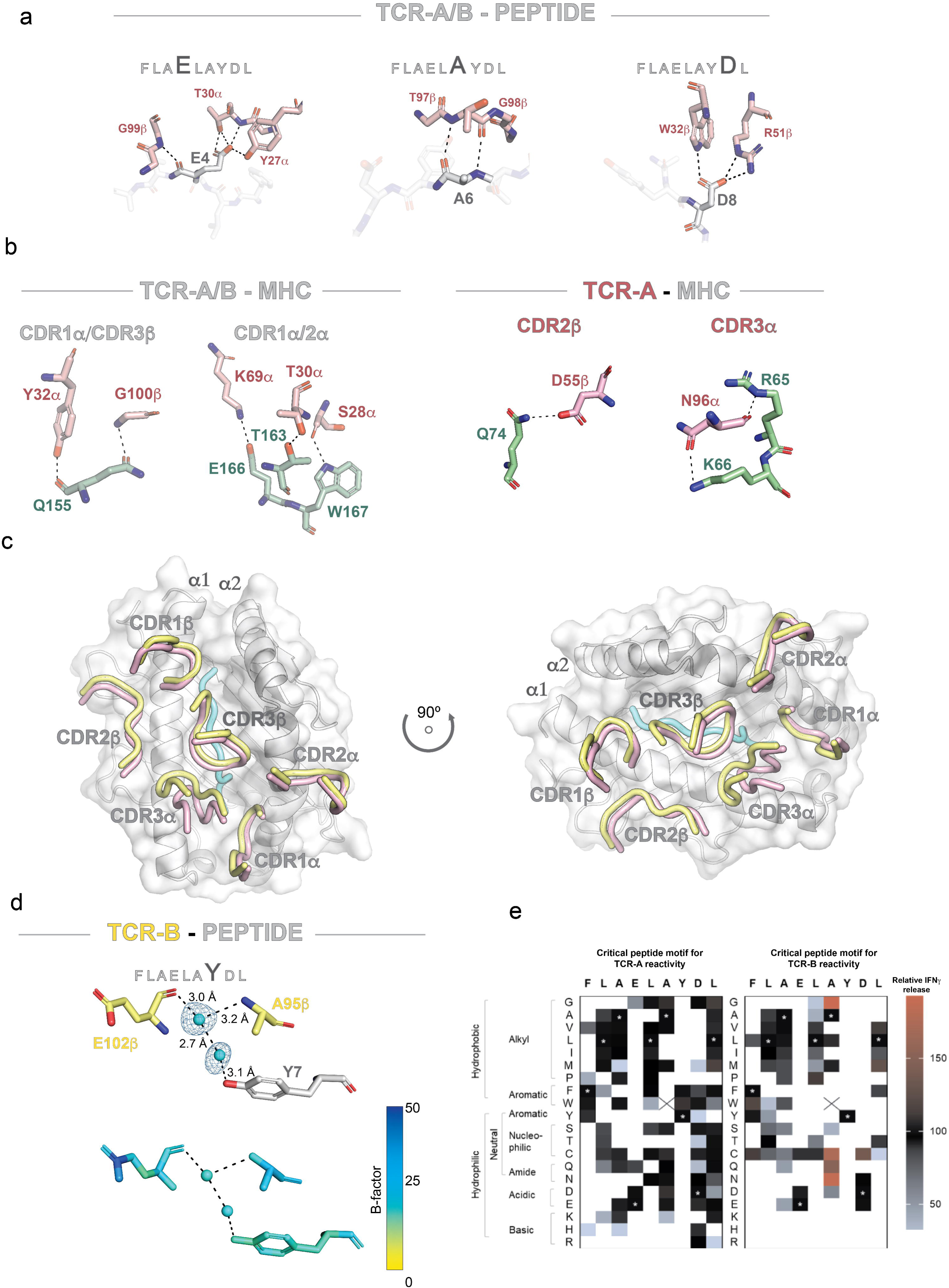
TCR:GPC3-9-mer and TCR:HLA-A2 interactions reveal fine-tuned differences between TCR-A and TCR-B. a, Close-up details of TCR-A/B polar interactions with the 9-mer peptide. Peptide and TCR residues are displayed in grey and pink color, respectively. Dashed lines depict hydrogen bonds or other polar/electrostatic interactions such as salt bridges (TCR R50β:peptide D8). The TCR CDR loops are labeled accordingly. **b,** TCR:HLA-A2 contacts conserved in both TCR-A and TCR-B (left panel). Interactions that are TCR-A-specific are also shown (right panel). HLA-A2 residues are highlighted in green color. **c,** Two views of the structural alignment of TCR-A and TCR-B CDR loops over the pMHC complex, represented as a grey surface with embedded cartoon. The 9-mer peptide backbone is displayed in cyan color, whereas the TCR CDRs are pink and yellow. **d,** Top panel, water-mediated contacts established by peptide Y7. Water molecules are represented as cyan spheres, along with their 2Fo-Fc electron density signals at a contour level of 1 sigma, shown as a thin blue mesh. The contacts formed between atom pairs (dashed lines) as well as their distances in Angstroms are shown. Bottom panel, the water-bridge snapshot is colored by B-factor values. **e,** X-scan mutational heat map for TCR-A and TCR-B recognition of GPC3-9-mer. Every position in the 9-mer peptide was replaced by alternative residues, and the degree of T-cell activation was determined by means of IFNγ release.

There are also conserved TCR interactions that involve residues on both helices of the MHC platform (Fig. 3b and Supplementary Tables 4, 5, 9 and 10). TCR-A, however, forms unique interactions with the MHC via the CDR2β and CDR3α loops, suggesting that both TCRs share a global binding footprint yet a non-identical framework of contacts to target the same antigen (Fig. 3b and Supplementary Tables 4,5). Structural superposition of the ternary complexes on the MHC molecules captures how the TCR-A and TCR-B CDR loops settled over the pMHC surface, involving a large occupancy area and leaving only the α2 N-terminal helix free of engagement (Fig. 3c).

Overall, these data describe equivalent docking geometries at a global level but distinct local engagement of the HLA-A2:GPC3-9-mer complex.

Likewise, the fine chemistry associated with engagement of the 9-mer peptide by TCR-A and TCR-B is not identical. In particular, the crystal structure with bound TCR-B shows additional contacts mediated by water molecules that involve the 9-mer Y7 residue (Fig. 3d). More specifically, the electron density and local and surrounding B-factors support the presence of a two well-ordered water bridge that connects the peptide antigen with the TCR-B CDR3β loop (Fig. 3d). This feature is not observed in the TCR-A complex. Since TCR-A and -B differ only in the sequence of CDR3α, we explored whether this difference underlies the distinct water-mediated contact observed for the TCR-B complex at peptide Y7. Structural alignment of both TCR:pMHC complexes relative to the MHC α1-α2 platform revealed that the different positioning of the CDR3α loop at the TCR:pMHC interface propagates into a subtle displacement of the CDR3β loop near the peptide Y7 side chain (Supplementary Figure 3). We hypothesize that this CDR3α-mediated shift of CDR3β contributes to the specific local hydration pattern that favors water-bridge formation in the TCR-B complex but not in its TCR-A counterpart.

### X-scan analysis supports a key role for GPC3-9-mer Tyr7

To assess the relevance of each peptide residue in its recognition by TCR-A and TCR-B, we mapped the tolerance to non-canonical peptide sequence variants through an X-scan screen (Fig. 3e and Supplementary Figure 4). In general, most of the residues shared by the two TCRs were located at the anchoring sites (positions 2 and 9) and position 3 (P3). P1 and P9 were highly permissive for TCR-A. Within MHC anchor sites, P4 and P7 were very restrictive in the two TCRs, accepting only a few amino acids, some of them being unique to each TCR. TCR-B was also very conservative and tolerated a scarce number of amino acids at P8, while TCR-A showed high permissiveness, accepting up to 13 different amino acids, most of them unique for this TCR. Regarding P5 and P6, TCR-A and B showed relatively high permissiveness, sharing many amino acids. Most of the common amino acids had alkyl or neutral side chains and were small/medium in size. This high similarity in terms of amino acids tolerated at P5 and P6 between TCR-A and TCR-B could be related to the fact that both TCRs share the same TCRβ chain. Interestingly, some amino acids at P6, such as G, N, C and Q, enhanced the reactivity of TCR-B, compared to the wild-type residue (A). Importantly, and consistent with the structural studies and binding assays, we found that replacement of Y7 was not tolerated by any amino acid in the case of TCR-B, indicating that the Y side chain oxygen plays an important role in antigen recognition (Fig. 3e). Opposite to this, position 7 tolerated either F or W when the test was performed with TCR-A, thus suggesting a specific role for the Y7 hydroxyl in TCR-B recognition, consistent with the water-bridge network described above. In summary, the TCR hot spots on the FLAELAYDL peptide are P4 and P7 for TCR-A, and P4, P7 and P8 for TCR-B, with P7-Y being irreplaceable.

Overall, the mutational and structural analyses demonstrate that TCR-A and TCR-B recognize highly similar antigenic footprints, with only subtle local differences in the chemistry of the binding interface. These observations argue against major differences in epitope specificity as the basis for their distinct therapeutic activities, and instead suggest that the functional divergence arises from more subtle features of productive antigen engagement.

### The GPC3**-**11-mer peptide conformation within HLA-A2 prevents TCR recognition

Previous studies suggested that an extended version of the 9-mer peptide with two additional C-terminal amino acids (FLAELAYDLDV) could also drive or potentiate TCR recognition. Having shown that this GPC3-11-mer adopts a distinct, protruding conformation within the HLA-A2 groove (Fig. 1), we next asked whether this altered topology is compatible with TCR engagement. We structurally aligned the HLA-A2:GPC3-11-mer complex with the ternary TCR-B:HLA-A2:GPC3-9-mer complex. Compared with the binding interface generated by the 9-mer peptide, the 11-mer peptide protrudes in a manner such that, considering a similar docking mode, it would produce severe steric clashes with the TCR CDRs, thus preventing coupling of the TCR-pMHC complex (Fig. 4a). Therefore, while both 9- and 11-mer peptides lead to productive pMHC folding states, the longer 11-mer version may not be compatible with TCR recognition. We also performed binding studies with TCR-A and TCR-B with the 11-mer-refolded HLA-A2 complex, but observed undetectable association in both cases (Fig. 4b). Parallel crystallization trials with the TCR-B and the MHC loaded with the 11-mer peptide failed to yield crystals, further supporting the lack of a plausible TCR-pMHC interaction.

**Fig. 4:**
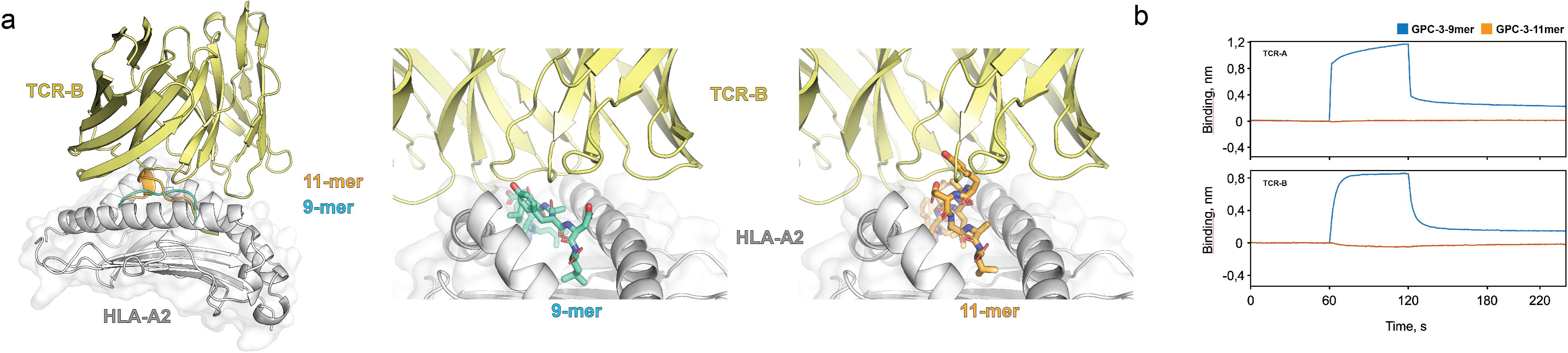
The 11-mer peptide conformation within the HLA-A2 does not enable TCR binding. a, Left panel, structural overlay, relative to the MHC molecule, of HLA-A2:GPC3-11-mer and the ternary TCR-B:HLA-A2:GPC3-9-mer complex. The 9- and 11-mer peptide backbones are shown in cyan and orange color, respectively, within the HLA-A2 groove, displayed in grey. TCR-B is represented as yellow cartoon. Middle and right panels, enhanced detail of the 9- and 11-mer peptides and their configuration at the TCR-B interface, with the peptide residues highlighted as sticks. The TCR-B:HLA-A2:GPC3-9-mer image was obtained from the atomic coordinates of the crystal structure, while the TCR-B:HLA-A2:GPC3-11-mer representation is the result of the structural alignment described in **a**. **b,** Real-time association and dissociation curves monitored through biolayer interferometry for the binding of refolded HLA-A2:GPC3-9-mer and HLA-A2:GPC3-11-mer complexes to surface-captured TCR-A or TCR-B.

We conclude that TCR recognition is defined by the peptide sequence on the one hand, and by the peptide folding mode within HLA-A2 on the other hand. We also hypothesize that the 11-mer peptide may have been trimmed by proteases (cell culture serum proteases or cell proteases) in previous functional assays^12^, resulting in ultimate presentation of trimmed 9-mer peptide versions, consistent with our structural and binding studies.

### Functional divergence emerges during target-cell engagement

Since neither structural analyses nor soluble binding measurements adequately explained the distinct therapeutic performance of TCR-A and TCR-B, we next investigated whether functional differences first arise during the earliest stages of T cell–tumor cell interactions.

Primary human CD8⁺ T cells expressing either TCR-A or TCR-B were co-cultured with GPC3-positive HepG2 cells, and conjugate formation was quantified over time by flow cytometry (Fig. 5a,b). The GPC3-negative EO771 cell line served as an antigen-negative control to assess nonspecific cell-cell interactions (Supplementary Fig. 5).

**Fig. 5:**
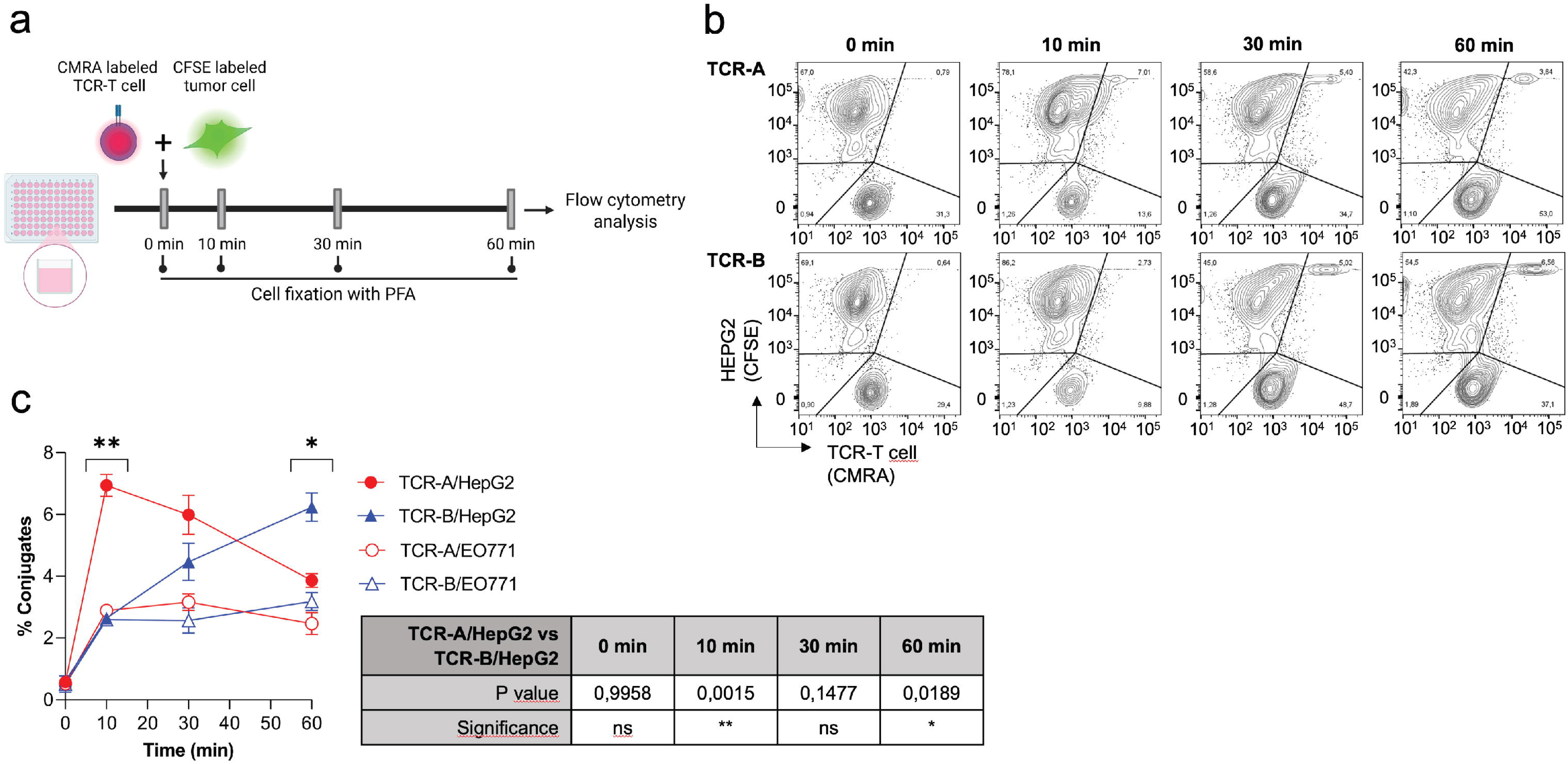
Distinct kinetics translate into different residence times in conjugate formation assays. a, Experimental setup: TCR-A and TCR-B T cells were labeled with CMRA, and tumor cell lines were stained with CFSE. Co-cultures were established at a 1:1 effector-to-target (E:T) ratio and fixed with paraformaldehyde (PFA) at the indicated time points. TCR-T cells and tumor cells seeded alone served as negative controls. **b,** Representative flow cytometry plots illustrating conjugate formation between HepG2 tumor cells and TCR-A or TCR-B T cells, identified as CFSE+ CMRA+ double-positive populations. **c,** Percentage of conjugates over time. Data are represented as mean ± SD (N=3 replicates per co-culture). Statistical significance was determined using a two-way ANOVA. *p < 0.05; **p < 0.01.

Both TCR-engineered T-cell populations formed antigen-dependent conjugates with HepG2 cells, but their temporal dynamics differed markedly (Fig. 5c). TCR-A T cells rapidly established target-cell conjugates and exhibited significantly higher frequencies of double-positive events during the early stages of the assay. However, this advantage progressively diminished over time. In contrast, TCR-B T cells displayed slower initial engagement but maintained stable interactions throughout the assay, ultimately surpassing TCR-A at the latest time point examined (Fig. 5c).

Comparison with the antigen-negative EO771 control further supported these distinct interaction dynamics. TCR-A maintained significantly higher conjugate formation with HepG2 cells throughout most of the assay, whereas TCR-B only became significantly different from background at the final time point (Supplementary Table 15). These observations indicate that TCR-A promotes rapid but relatively transient target-cell interactions, whereas TCR-B favors the progressive establishment of more durable cell-cell contacts.

Collectively, these results identify target-cell engagement as the earliest stage at which the functional behavior of the two receptors begins to diverge. The TCRs differ in the temporal stability of their interactions with target cells, suggesting that prolonged conjugate persistence may contribute to the superior therapeutic performance of TCR-B.

### TCR-B T cells display superior persistence compared with TCR-A T cells in repetitive antigen challenge (RAC) assays

The distinct target-cell engagement dynamics observed in conjugate assays prompted us to determine whether these differences influenced T-cell performance under sustained antigen exposure. To mimic the repetitive stimulation experienced by engineered T cells within solid tumors, we established a repetitive antigen challenge (RAC) assay in which TCR-engineered T cells were sequentially exposed to fresh GPC3-positive HepG2-LUC-GFP target cells every 24 hours for five consecutive rounds (Fig. 6a,b).

**Fig. 6:**
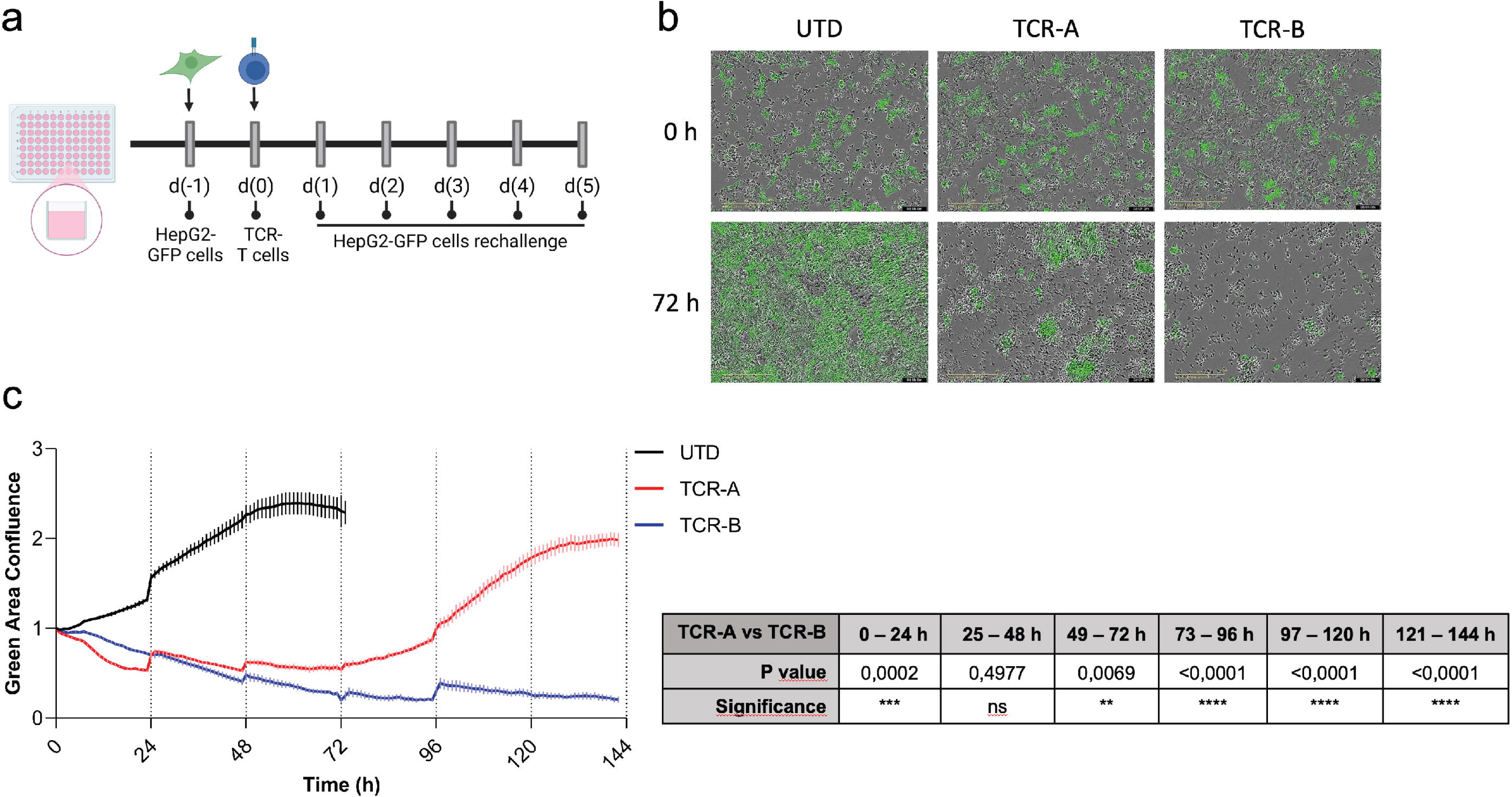
TCR-B T cells display superior persistence compared with TCR-A T cells in repetitive antigen challenge (RAC) assays. a, HepG2-LUC-GFP tumor cells were seeded in 96-well plates; TCR-A or TCR-B T cells were added the following day at a 3:1 (E:T) ratio. New challenges with HepG2-LUC-GFP cells were performed every 24 h for a total of five consecutive rounds. Untransduced T cells (UTD) served as a negative control for the first two challenges. **b**, Representative Incucyte images of the indicated conditions and time points. **c,** Green area confluence over time. Data are represented as mean ± SD (N=5 independent co-cultures). Statistical significance was determined using a two-way ANOVA of the area under the curve (AUC) for each condition across the indicated time intervals. ns, not significant; **p < 0.01; ***p < 0.001; ****p < 0.0001.

During the initial challenge, TCR-A T cells exhibited significantly faster tumor-cell elimination than TCR-B T cells, consistent with their more rapid target-cell engagement observed in the conjugate assay (Fig. 6c). However, this early advantage progressively disappeared. Following the second antigen challenge, the cytotoxic activities of both receptors converged, after which their functional trajectories diverged markedly.

Whereas TCR-A progressively lost cytotoxic activity with successive rounds of antigen exposure, TCR-B maintained efficient tumor-cell control throughout the entire experiment. By the fourth and fifth challenges, TCR-A was no longer capable of effectively controlling tumor growth, while TCR-B preserved robust and sustained cytotoxic activity (Fig. 6c). Thus, although TCR-B displayed slower initial tumor-cell killing, it consistently outperformed TCR-A under conditions of chronic antigen stimulation.

These observations demonstrate that the principal functional difference between the two receptors does not lie in their initial cytotoxic potential but rather in their ability to maintain effector function during prolonged antigen exposure. Importantly, this divergence only becomes evident under repetitive stimulation, a setting that more closely resembles the continuous antigenic pressure encountered within solid tumors.

### Enhanced functional endurance translates into superior antitumor efficacy in vivo

Finally, we asked whether the superior functional endurance displayed by TCR-B under repetitive antigen stimulation translated into improved therapeutic efficacy in vivo. NSG mice bearing established PLC/PRF/5-A2 hepatocellular carcinoma xenografts received a single adoptive transfer of TCR-A-, TCR-B-, or untransduced (UTD) CD8⁺ T cells, followed by systemic IL-2 administration (Fig. 7a).

**Fig. 7:**
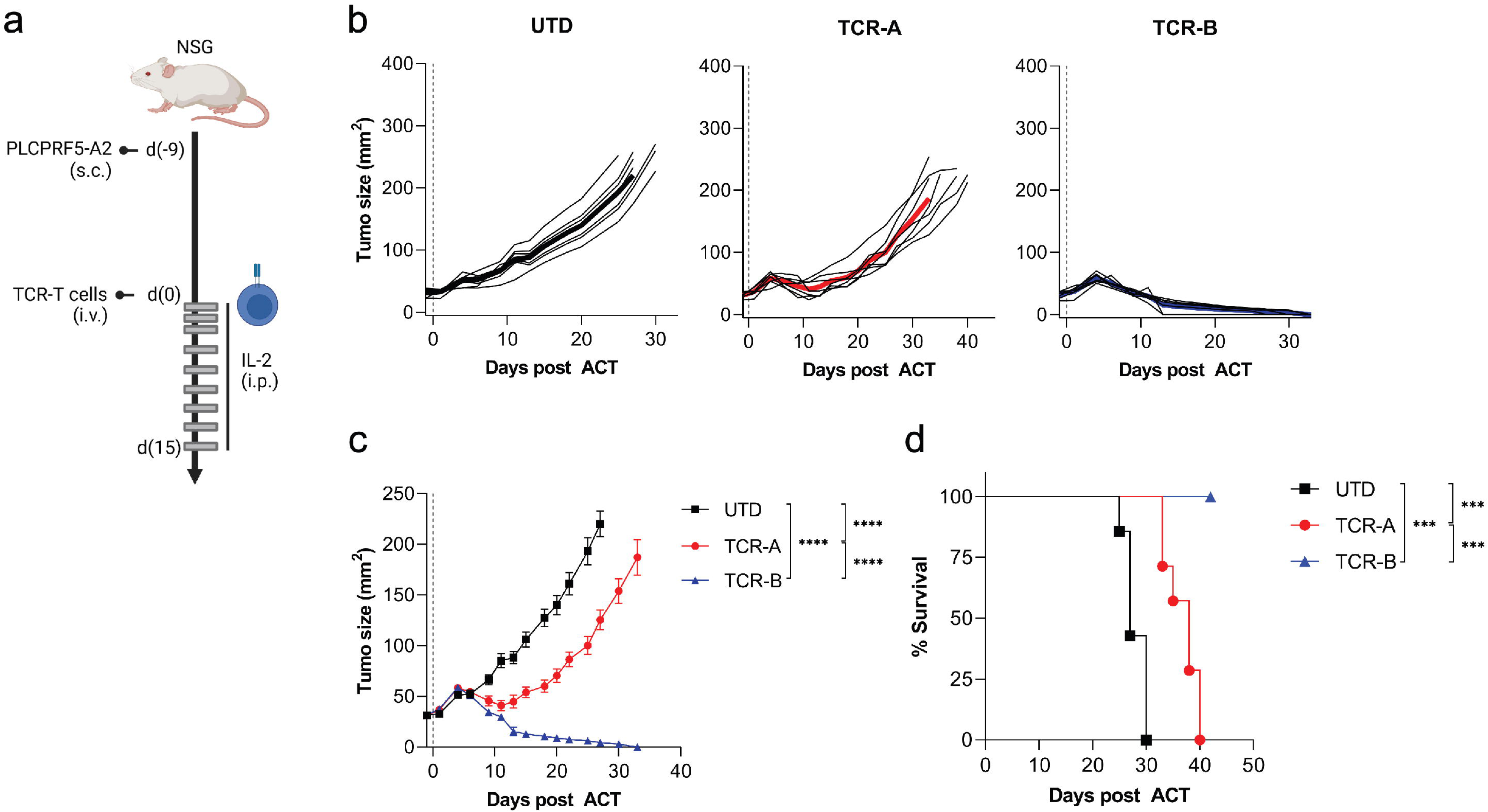
TCR-B T cells outperform TCR-A T cells in ACT assays. a, Six- to eight-week-old NSG mice were subcutaneously (s.c.) implanted in the right flank with PLC/PRF/5-A2 cells. Nine days post-implantation, mice were administered 7-8 × 10^6^ TCR-A or TCR-B T cells via intravenous (i.v.) injection (N=7 mice/group). The total number of mTCRβ+ CD8+ T cells was equalized between groups by supplementing with autologous UTD CD8+ T cells. The control group received an equivalent number of UTD CD8+ T cells (N=7 mice). Recombinant human IL-2 was administered intraperitoneally (i.p.) on days 1, 2, 3, 5, 7, 9, 11, 13, and 15, and s.c. adjacent to the tumor site on days 3, 5, 7, 9, 11, 13, and 15 post-ACT. **b**,**c**, Tumor size (mm^2^) following ACT. **b**, Individual tumor growth curves for each mouse. **c,** Average tumor size per group. Data are represented as mean ± SEM. **d**, Overall survival. Statistical significance was determined using non-linear fit test (curve fit) for **c** and the Mantel-Cox test for **d**. ***p < 0.001; ****p < 0.0001.

Both engineered T-cell products significantly delayed tumor progression compared with UTD controls, confirming that both receptors efficiently recognize the GPC3 antigen in vivo (Fig. 7b,c). However, their therapeutic outcomes diverged substantially. While TCR-A induced transient tumor control, tumors eventually progressed in all treated animals. In contrast, TCR-B mediated complete tumor eradication, leading to durable responses and long-term survival in every treated mouse (Fig. 7b–d).

These in vivo findings closely mirror the functional differences observed throughout the preceding experiments, with TCR-B consistently outperforming TCR-A under conditions that require sustained antigen engagement. The superior tumor control achieved by TCR-B therefore appears to reflect its enhanced capacity to maintain productive effector function during continuous antigen exposure rather than differences in receptor affinity.

Collectively, the structural, biophysical, cellular and in vivo data support a unified model in which productive antigen engagement and sustained functional endurance, rather than soluble affinity alone, are the principal determinants of therapeutic performance for GPC3-specific TCR-T cells.

## Discussion

We investigated two TCRs with different antitumor efficacies despite targeting the same HLA-A*02:01-restricted GPC3(522-530) 9-mer peptide. While both TCR-A and TCR-B establish highly analogous contacts with HLA-A2:GPC3-9-mer, TCR-A drives faster tumor-cell engagement and early cytotoxicity, whereas TCR-B promotes greater stability of target-cell conjugates, superior properties under repeated antigen challenge, and enhanced antitumor efficacy in vivo. These results support the conclusion that, in solid-tumor adoptive T-cell therapy, therapeutic performance is not predicted by antigen specificity or affinity between soluble components alone.

We provide several high-resolution crystal structures showing that HLA-A2 can allocate not only different peptide lengths, but different conformations for a given peptide, as shown with GPC3-9-mer. We demonstrate that the 9-mer peptide can adopt different folds within the MHC groove. What is not yet conclusive is whether the HLA-A2:GPC3-9-mer complex exists in a mixed ratio of conformations in solution, or whether TCR proximity induces a pre-docking rearrangement of the peptide antigen that favors complete TCR-pMHC assembly. The 11-mer GPC3(522-532) peptide, however, binds efficiently to HLA-A2, but in a manner such that it adopts an alternative fold, accompanied by a severe upward protrusion outside HLA-A2’s groove that prevents TCR docking. Unlike with the 9-mer peptide, we could not detect a productive binding through biolayer interferometry to the HLA-A2:GPC3-11-mer complex. Neither could we obtain crystals of ternary TCR-pMHC complexes when the MHC was loaded with the 11-mer peptide. We conclude that these GPC3-specific TCRs target the 9-mer version. Additionally, these findings indicate that previous activity associated to the 11-mer peptide^12^ may be attributable to protease-mediated trimming into the shorter antigenic species. These assays were performed in T2 cells^12^. While these cells lack the transporter associated with antigen processing (TAP), the cells were cultured in the presence of 10% fetal bovine serum, which could be a source of proteases leading to trimming of the longer peptide.

When aligned over the MHC platform, the unbound and TCR-bound HLA-A2:GPC3-9-mer structures reveal prominently distinct arrangements of peptide Leu5 within the MHC groove. Relative to its counterpart in the TCR-unbound structure, Leu5 is rotated in a manner such that its backbone and side chain shift their orientation, creating a gateway for CDR3β and subsequent TCR docking. This observation suggests that, apart from length, conformational permissiveness within HLA-A2 is central to GPC3-peptide recognition.

While both TCRs recognize the same GPC3-9-mer peptide through nearly identical binding footprints dictated primarily by peptide residues E4, A6 and D8, crystal structures reveal subtle differences in intermolecular contacts. More specifically, the TCR-B CDR3β loop associates with peptide Y7 - which resulted critical for antigen recognition in a mutational X-scan assay - through a water bridge involving two well-ordered water molecules. This specific contact is not recapitulated in the ternary TCR-A:pMHC structure, where, as judged by the X-scan analysis, position 7 in the peptide tolerates replacement with any other aromatic amino acid, i.e., F and W. This observation raises the possibility that, unlike the TCR-A, the TCR-B αβ pairing imposes a fine-tuned and distinct geometric constraint at the TCR-B:HLA-A2:GPC3-9-mer interface involving the Y7 side chain oxygen of GPC3-9-mer.

Soluble binding analyses confirmed specific recognition of GPC3 antigen for both TCRs, with TCR-B featuring slower on and off-rates. While the structural and biophysical data support specific engagement of HLA-A2:GPC3-9-mer - with TCR-B showing a more stable TCR-pMHC interaction based on the slower off-rate - there is no direct evidence that could be inferred from these data for the superior therapeutic efficacy observed for TCR-B. Together, our findings reinforce the growing concept that productive TCR signaling cannot be inferred solely from measurements of soluble affinity or dwell time. Instead, productive antigen engagement likely emerges from a complex integration of structural compatibility, receptor dynamics and mechanical interactions occurring at the immunological synapse.

Whereas molecular analyses revealed subtle differences between the two receptors, functional assays uncovered progressively increasing divergence as antigen exposure became sustained. TCR-A produced early target-cell engagements, whereas TCR-B led to more persistent assemblies at later stages. Particularly relevant is the fact that this divergence is associated with functional endurance. TCR-A presented potent cytotoxicity at initial time points, but activity dropped with time under successive antigen challenge. On the contrary, TCR-B preserved its antitumor properties throughout the global antigen challenge.

The therapeutic translation of this discrepancy was confirmed in the in vivo ACT model. While TCR-A was able to control tumor progression, it was only TCR-B that completely ablated the tumor and promoted enhanced survival.

These findings suggest that therapeutic TCRs should not necessarily be viewed as receptors that maximize immediate cytotoxicity, but rather as receptors capable of preserving productive effector function during continuous antigen stimulation. This distinction is particularly relevant in solid tumors, where engineered T cells experience persistent antigen exposure over prolonged periods rather than isolated encounters with target cells.

Conventional short-term cytotoxicity assays primarily measure the initial activation potential of engineered lymphocytes, whereas serial challenge more faithfully reproduces the chronic stimulation encountered within the tumor microenvironment. Under these conditions, TCR-B consistently maintained tumor control while TCR-A progressively lost functional capacity, suggesting that functional endurance represents a previously underappreciated property of therapeutically effective TCRs.

Previous studies by other groups support these findings. Vazquez-Lombardi and coworkers reported discrepancies between TCR binding and cellular activation^15^. The authors propose that functional screening supports the identification of TCR variants with enhanced potency. This is relevant because a tighter engagement may result counterproductive, as demonstrated in previous TCR engineering studies. Sibener and colleagues also showed that structural determinants of the TCR:pMHC interface can uncouple binding affinity from signaling performance^16^. Therefore, the superior properties observed for TCR-B are best described as derived from productive antigen engagement and enhanced functional endurance, rather than as a direct result of higher affinity measured with soluble reagents.

In conclusion, our study establishes a mechanistic framework linking antigen conformational permissiveness, productive molecular recognition and sustained functional endurance to therapeutic efficacy. These findings encourage the implementation of metrics regarding functional endurance into TCR selection workflows for adoptive T cell therapy. In the context of solid tumors, where engineered T cells face sustained stimulation through successive antigen challenge, metrics related to serial cytotoxicity, cellular endurance upon repetitive antigen challenge, stability of target-cell conjugates and specificity towards the tumor antigen may represent a stronger predictive model of therapeutic performance than the affinity measured with soluble TCR and pMHC reagents.

## Methods

### Mice

NOD.Cg-PrkdcscidIL2rgtm1Wjl/SzJ (NOD scid gamma, NSG) mice were purchased from Jackson Laboratory. Mice were bred and housed under specific pathogen-free conditions in the CIMA animal facility. Animal care and experiments were conducted in accordance with the ARRIVE guidelines (Animals in Research: Reporting In Vivo Experiments) and approved by our institutional ethics committee (048-21), in compliance with Spanish regulations.

### Peptides, plasmids and retroviral vectors

The GPC3-9-mer FLAELAYDL and GPC3-11-mer FLAELAYDLDV were synthetized by Gencust (purity > 95 %). The pcDNA3.4 plasmid was purchased from bioNova. The pET28 plasmid was kindly provided by Dr. Lasa. The pMD2.G plasmid was obtained from the Addgene repository (provided by Didier Trono; Addgene plasmid #12259). The Murine Stem Cells Virus (MSGV1)-Mu(*C) Acceptor plasmid was created as previously described^12^.

### Cell lines

The HepG2 tumor cell line (CVCL_0027, pediatric hepatocellular carcinoma) was authenticated by short-tandem repeat (STR) profiling (AmpFLSTR Identifiler Plus PCR Amplification Kit) in 2023. Cells were transduced with the MSCV-LUC-IRES-GFP vector (Addgene plasmid # 20672) to generate the HepG2-LUC-GFP cell line. Tumor cells were cultured in “Tumor medium” [MEM-GlutaMAX (GIBCO), 10% Fetal Bovine Serum (FBS) (SIGMA), 1% sodium pyruvate (GIBCO), 1% non-essential amino acids (GIBCO), 10 mM HEPES (GIBCO), 100 U/mL Penicillin/Streptomycin (P/S) (GIBCO), and 10 μg/mL gentamicin (GIBCO)].

Additional cell lines used in this study include: CHO-S cells: a CHO cell line optimized for recombinant protein production, purchased from Invitrogen (2023), and cultured in ExpiCHO^TM^ Expression Medium (Thermo Fisher). Platinum-A (PLAT-A) cells: a retroviral packaging cell line, purchased from Cell Biolabs (2018), and cultured in “PLAT-A medium” [DMEM-GlutaMAX (GIBCO), 10% FBS, 1% sodium pyruvate, 1% non-essential amino acids, 10 mM HEPES, 100 U/mL P/S and 10 µg/mL gentamicin] supplemented with Puromycin (1 μg/mL) and Blasticidin (10 μg/mL). T2 cells: TAP-deficient HLA-A2+ cells (CVCL_2211), purchased from ATCC (2017) and cultured in “complete (c)RPMI medium” [RPMI-1640-GlutaMAX (GIBCO), 10% FBS, 20 mM of HEPES, 100 U/mL of P/S, 10 µg/mL gentamicin, and 50 µM 2-mercaptoethanol (GIBCO)]. PLC/PRF/5-A2 cells: a human hepatocellular carcinoma cell line (CVCL_0485), authenticated by STR profiling and cultured in “Tumor medium”. EO771 cells: a murine epithelial-like mammary carcinoma cell line, purchased from ATCC (2023) and cultured in “complete DMEM medium” [DMEM (ATCC), 10% FBS and 20 mM HEPES].

### Production and purification of the recombinant TCRs and pMHC

The gene sequences encoding the TCR-B alpha and beta chains were synthesized by GeneUniversal. The alpha chain was designed to incorporate N164Q, N198Q and N209Q substitutions, a BamHI restriction site between the variable and constant regions, and a C-terminal 6xHis tag. The beta chain construct included an N203Q substitution and a XhoI restriction site located between the variable and constant regions. The sequences were provided in pUC57 vectors flanked by EcoRI and HindIII.

The gene sequence encoding the TCR-A alpha chain variable region was synthesized by GeneUniversal (the beta chain is common to both TCRs). The sequence was provided in pUC57 vector flanked by EcoRI and BamHI restriction sites and was digested for 15 minutes at 37 °C using FastDigest Restriction Enzymes (Thermo Scientific). As the constant region is common to both TCRs, the previously obtained TCR-B alpha chain construct was digested with EcoRI and BamHI, and the resulting fragments from both digestions were ligated using T4 DNA ligase (Thermo Scientific).

The cloning, production and purification of the recombinant TCRs and pMHC complexes was performed as previously described^18^. Briefly, genes encoding the alpha and beta chains of TCR-A and TCR-B were cloned into pcDNA3.4 expression vectors. The plasmids were separately co-transfected into CHO-S cells, and the secreted recombinant TCRs were harvested from the culture supernatants eight days post-transfection. Following dialysis and filtration, the TCRs were purified via HisTrap affinity chromatography and Superdex 200 size exclusion chromatography. Concurrently, HLA-A*02:01 and β2 microglobulin genes were cloned into pET28 vectors and expressed as inclusion bodies in *E. coli* BL21(DE3) (Agilent) cells upon IPTG induction. The isolated inclusion bodies were solubilized in 8.0 M urea, refolded by dilution in a buffer containing synthetic GPC3-9-mer, GPC3-11-mer, or PR3(169-177) peptides, dialyzed for 72 hours, and purified using HiTrap Q FF anion exchange column. Finally, HLA-A2:GPC3 complexes were concentrated to 5 mg/mL, and ternary complexes were formed by mixing stoichiometric amounts of the HLA-A2:GPC3 complexes with either TCR-A or TCR-B, and subsequently concentrated to 5 mg/mL.

### Crystallization of the binary pMHC and ternary TCR:pMHC complexes

Each complex was screened against more than 500 crystallization conditions using the sitting drop vapor diffusion method and 20 °C incubation. A Crystal Gryphon LCP platform (Dunn Labortechnik GmbH) was used for these trials. Drop volumes were set at 300 nanoliters for the crystallization reagent in each condition, while the protein sample drop volumes ranged from 250 to 300 nanoliters. Crystals of HLA-A2:GPC3-9-mer appeared in 24-48 hours and were obtained in 0.2 M NaF, 0.1 M BIS-Tris propane pH 7.5, and 20% w/v PEG 3350. Crystals of HLA-A2:GPC3-11-mer appeared in 1-2 weeks and were obtained in 0.005 M MgCl2, 0.005 M NiCl2, 0.005 M CoCl2, 0.005 M CdCl2, 0.1 M HEPES pH 7.0, and 12% w/v PEG 3350. Crystals of TCR-A:HLA-A2:GPC3-9-mer appeared in 24-48 hours and were optimized in 0.1 M MIB buffer pH 8.0 and 23% w/v PEG 1500. Finally, crystals of TCR-B:HLA-A2:GPC3-9-mer appeared in 24-48 hours and were obtained in 0.2 M NaCl, 0.1 M MES, and 20% PEG 6000. Crystals were harvested, soaked in crystallization medium supplemented with 20% glycerol (TCR-B:HLA-A2:GPC3-9-mer) or 20% ethylene glycol (HLA-A2:GPC3-9-mer, HLA-A2:GPC3-11-mer and TCR-A:HLA-A2:GPC3-9-mer), and cryo-cooled in liquid nitrogen before diffraction analyses.

### Diffraction analysis, data processing and structure determination and refinement

X-ray diffraction and data collection were performed on the BL13-Xaloc beamline at the Alba synchrotron facility (Cerdanyola del Vallès, Barcelona, Spain). Diffraction datasets were collected for all crystallized complexes. The diffraction data was reduced and integrated with XDS^19^ and then merged and scaled with *AIMLESS*^20^ in the CCP4 Suite^21^. 5% of reflections were excluded for validation purposes during the refinement process. The structures were solved by molecular replacement *via* Phaser^22^. For HLA-A2:GPC3 complexes, the two independent templates used for phasing were PDB IDs 7KGP and 7EJL, corresponding to HLA-A*02:01 and β2 microglobulin, respectively. For the ternary complexes, the same two templates were used for the pMHC, and three additional templates were used for the TCRs: 7Z50 for the constant domains, and 3TJH and 8RO5 for the variable domains of alpha and beta chains, respectively. Molecular replacement solutions were subsequently refined with Refmac5^23^ or phenix.refine^24^. The final models were generated after iterative cycles of manual building in Coot^25^ and further refinement prior to data deposition in the Protein Data Bank.

### Kinetic characterization using biolayer interferometry

The affinities of TCR-A and TCR-B for the pMHC complexes were tested by biolayer interferometry using a BLItz system (Sartorius). His-tagged TCRs were immobilized onto Ni-NTA-coated biosensors (Sartorius) before association of increasing concentrations (15, 30 and 60 μM) of the pMHC complex for 60 s followed by dissociation using HEPES-buffered saline (HBS) pH 7.4 for 120 s. The obtained sensorgrams were corrected by substracting the signal obtained with HBS and fitted to a 1:1 Langmuir binding model using the BLItz software.

### Off-target study of GPC3-specific TCRs with X-scan assay

An X-scan assay was performed and adapted to our conditions. Briefly, a library of 172 peptides, in which each residue of the FLAELAYDL target peptide was sequentially mutated to all 19 other naturally occurring amino acids was synthesized by Innovative Peptide Solutions (Berlin, Germany) to a purity > 70%. Each peptide was analyzed by LC-MS. Capping of truncated peptides after each synthesis step was included to avoid the formation of deletion peptides and thus the detection of false-positive T-cell responses. X-scan assay was performed as follows. TCR-A and TCR-B-engineered CD8 T cells from non-HLA-A*02:01 donors were thawed and recovered in “T cell medium” [a 1:1 mixture of AIM-V (Invitrogen) and RPMI 1640-glutamax (GIBCO), containing 5% heat-inactivated human AB serum (SIGMA), 12.5 mM HEPES, 100 U/mL of P/S, and 10 μg/mL gentamicin] supplemented with IL-7 and IL-15 for 2 days before use. For each triplicate well of the 96-well plates, 5 × 10^3^ mTCRβ^+^ cells were cultured with 2 × 10^4^ T2 cells in the presence of each peptide at a final concentration equal to the EC_90_ of the index FLAELAYDL peptide (0.5 µM in the case of TCR-B and 0.05 µM in the case of TCR-A). After overnight co-culture, the media was collected and assayed for IFNγ by ELISA [human BD OptEIA Set (BD Biosciences)] according to the manufacturer’s instructions. The absorbance of IFNγ was then compared to the IFNγ release level after FLAELAYDL peptide stimulation and the ratios were calculated.

### Genetic modification of human T lymphocytes

The experimental protocol was performed as previously described^12^. Briefly, retroviral supernatants were generated by cotransfecting PLAT-A cells (Cell Biolabs) with TCR-A or TCR-B retroviral vectors (MSGV1) together with the pMD2.G helper plasmid. T cells isolated from the blood of healthy donors (Miltenyi) were activated (48 h) with Transact (Miltenyi) in T cell medium supplemented with IL-7 and IL-15. Activated T cells were then transduced with the retroviral particles encoding TCR-A or TCR-B. Transduction efficiency was assessed using FACS Canto II flow cytometer (BD Biosciences) based on the expression of murine (m) TCRβ (clone H57-597) and human CD8 (clones HIT8a, RPA-T8, and SK1) (cells were incubated with the mix of fluorophore-conjugated mAbs in FACS buffer (PEF Buffer + 0.05% azide) containing Beriglobin P (RT/15’)). To properly compare the different TCRs, the percentage of murine (m)TCRβ+ CD8 T cells in the different TCR-T cell lines was equalized by adding untransduced (UTD) T cells. The study was performed in accordance with the Declaration of Helsinki and Istanbul, and approved by the Ethics and Scientific Committee of the University of Navarra (2020.152). Written informed consent was obtained from all healthy donors.

### Rapid Expansion Protocol (REP)

TCR-A and TCR-B transduced T cells were cultured with 30 ng/mL OKT3 anti-CD3 antibody (BioLegend), 3000 U/mL of IL-2 (Proleukin) and irradiated feeder cells (40 Gy) (100:1 feeder:T-cell ratio) in T cell medium. After 3 days of culture, the medium was refreshed by adding 1/3 volume of T cell medium supplemented with 3000 U/mL of IL-2. On days 6 and 8 of the REP, half of the culture medium was removed and replaced with fresh T cell medium supplemented with 3000 U/mL of IL-2 at twice the remaining culture volume. The purity of the expanded cultures was assessed by flow cytometry by measuring the percentage of hCD8/mTCRβ double-positive T cells. After completion of the REP, T cells were used for adoptive cell transfer or co-culture assays.

### Fluorescent labeling of cells

Fluorescent dye-5-(and-6)-carboxyfluorescein diacetate, succinimidyl ester (CFSE) (SIGMA) was used to label HepG2-LUC-GFP and EO771 tumor cells, while 5-(and-6)-(((4-chloromethyl)benzoyl)amino)tetramethylrhodamine (Orange CMRA) (Invitrogen) was used to label TCR-A and TCR-B T cells.

For tumor cell labeling, 4.4 × 10^6^ cells of each tumor cell line were washed and resuspended in 440 µL of PBS. The cells were incubated with CFSE at a final concentration of 0.125 µM for 10 min at 37 °C, followed by addition of bovine serum (BS) (LINUS) to a final concentration of 10% (v/v). Cells were centrifuged at 2000 rpm for 6 min, resuspended in 500 µL of “Tumor medium” and kept on ice until the conjugate assay. In parallel, 2.2 × 10^6^ cells of each modified T cell population were washed and resuspended in 220 µL of “Tumor medium”. The cells were incubated with Orange CMRA at a final concentration of 0.5 µM for 20 min at 37 °C, followed by two washing steps with RPMI-1640 (GIBCO). Finally, cells were resuspended in 500 µL “Tumor medium” and kept on ice until the conjugate assay.

### Formation and measurement of conjugates

Conjugate formation was assessed at four different time points (0, 10, 30 and 60 min), using one 96-well cell culture plate per time point. Cells were derived from a single donor, and three replicates were analyzed.

For the 0 min time point, 0.1 × 10^6^ TCR-T cell were seeded and mixed with an equal number of the corresponding tumor cell line in each well. As negative controls, TCR-T cells and tumor cells were seeded alone in separate wells. Immediately after mixing, 100 µL/well of 4% paraformaldehyde (Thermo Fisher Scientific) at 4 °C were added to fix the cells and stabilize formed conjugates. The plate was centrifuged at 2000 rpm for 2 min, supernatants were discarded, and cells were resuspended in 180 µL of FACS buffer. For the remaining time points (10, 30 and 60 min), TCR-T cells and tumor cells were mixed as described above, plates were centrifuged at 2000 rpm for 1 min to promote cell contact, and incubated at 37 °C for the indicated times. At the end of each incubation, 100 µL/well of 4 % paraformaldehyde at 4 °C were added, plates were centrifuged, supernatants discarded, and cells resuspended in 180 µL of FACS buffer. All samples were kept at 4 °C protected from light until analysis by flow cytometry.

Conjugate formation was quantified using FACS Canto II flow cytometer. Free effector cells were stained red alone, free target cells were stained green alone, while conjugates were double-positive, staining both red and green. Data were analyzed using FlowJo software (Tree Star).

### Repetitive antigen challenge

The experimental setup was adapted from a previously described protocol^17^. Briefly, 48-72 h before the experiment, HepG2-LUC-GFP cells were treated with IFNγ (100 U/mL) (Immunotool) to enhance HLA expression and antigen presentation. On the day of the experiment, wells of a 96-well cell culture plate (Greiner CELLSTAR^®^) were treated with Poly-L-ornithine (Merck) for 1 h, after which 8000 cells/well of HepG2-LUC-GFP cells were seeded and incubated at 37 °C overnight. The next day, TCR-T cells with IL-2 (10 U/mL) were added using a 3:1 effector:target ratio and incubated for 24 h. Every 24 h, 8000 cells/well of HepG2-LUC-GFP were added to each well along with IL-2 (10 U/mL), for a total of five challenges. HepG2-LUC-GFP cell growth and killing were monitored over time and imaged using a real-time live-cell imaging system (Incucyte, Essen Biosciences). UTD T cells were used as a negative control.

The experiment was performed in “Tumor medium” using cells from a single donor, with five replicate measurements.

### Adoptive Cell Transfer of engineered T cells

Six to eight-week-old NSG mice were subcutaneously implanted in the right flank with 2.5 × 10^6^ PLC/PRF/5-A2 human hepatocellular carcinoma cells. Tumor dimensions were assessed by caliper measurement 8 days after implantation, and mice were randomized into different treatment groups to ensure comparable mean tumor sizes and standard deviations across cohorts. On day 9, mice were administered 7-8 × 10^6^ mTCRβ+ engineered CD8 T cells via intravenous injection using the retro-orbital route. To compare different TCR-T cells, the total number of CD8 T cells was equalized between groups by supplementing with autologous UTD CD8 T cells. The control group received an equivalent number of UTD CD8 T cells as the total number of TCR-engineered CD8 T cells. Recombinant human IL-2 was administered intraperitoneally at a dose of 4 × 10^4^ IU/mouse on days 1, 2, 3, 5, 7, 9, 11, 13, and 15, and subcutaneously adjacent to the tumor site at 2 × 10^4^ IU/mouse on days 3, 5, 7, 9, 11, 13, and 15 following ACT.

Tumor growth was monitored three times per week by measuring perpendicular diameters using digital calipers, and survival rates were also recorded. In compliance with institutional ethical guidelines (protocol 048-21), mice were euthanized upon meeting any of the following criteria: a mean tumor diameter [(longest diameter + shortest diameter)/2] reaching 13 mm, evidence of tumor ulceration or necrosis, and/or signs of physical impairment (such as impaired mobility, lethargy, reduced physical activity, and marked weight loss).

### Statistical analysis

Statistical tests were performed using GraphPad Prism (v.8.4.0). To measure the dispersion of individual observations around the mean, the standard deviation (SD) was employed. This metric was utilized for analytical assays involving triplicate to quintuplicate determinations from a single representative experiment. In contrast, the standard error of the mean (SEM) served to estimate variability between group means. SEM was specifically used for biological experiments where data points were derived from different animals in each comparison group. Statistical evaluation of multiple comparisons was performed via two-way ANOVA. For the RAC assay, the area under the curve (AUC) was calculated for each interval, and the resulting values were analyzed using two-way ANOVA. Non-linear fit test was utilized to compare tumor growth curves, while the Mantel-Cox test was applied to analyze differential survival. In all cases, statistical significance was defined as p < 0.05.

## Acknowledgements

We are grateful to the staff of Xaloc beamline at ALBA Synchrotron for their assistance with X-ray diffraction data collection.

## Autorship contributions

Conceived research: SH and JLS; Performed experiments: LOO, GDR, EE, JLS; Data collection and analysis: LOO, JLS, SH; Draft writing: JLS, LOO and SH.

## Disclosure of conflicts of interest

The authors declare no competing interests.

## Data availability

Atomic coordinates and structure factors have been deposited and are available at the Protein Data Bank under the accession code PDB ID 9TM0 for HLA-A2:GPC3-9-mer, 9TMM for TCR-A:HLA-A2:GPC3-9-mer, 9TML for TCR-B:HLA-A2:GPC3-9-mer and 31CS for HLA-A2:GPC3-11-mer.

## Funding

This work was funded by Gobierno de Navarra Ayudas para la realización de Proyectos Estratégicos de I+D 2023-2026 (0011-1411-2023-000072, PITAGORAS) and Doctorados Industriales, Grant 0011-1408-2025-000017. Funding from the Ministry of Science, Innovation and Universities of Spain, grant PID2022-139888NB-I00/AEI/10.13039/501100011033/FEDER, UE, also supported this work.

## Notes

### Competing Interest Statement

The authors have declared no competing interest.

