## Supplementary Material for "Structure-function profiling identifies determinants of TCR-T cell therapeutic efficacy"

<sup>1</sup> Unit of Protein Crystallography and Structural Immunology, Navarrabiomed, Navarra, Spain.

<sup>2</sup> Public University of Navarra (UPNA), Pamplona, Navarra, Spain.

<sup>3</sup> Navarra Hospital Complex (CHN), Pamplona, Navarra, Spain.

<sup>4</sup> Instituto de Investigación Sanitaria de Navarra (IdiSNA), Pamplona, Spain.

<sup>5</sup> Program of Immunology and Immunotherapy, CIMA-University of Navarra, Pamplona, Spain.

<sup>6</sup> Centro de Investigación Biomédica en Red de Enfermedades Oncológicas, Instituto de Salud Carlos III, Madrid, Spain.

\*To whom correspondence should be addressed:

### SUPPLEMENTARY MATERIALS.

**Supplementary Figure 1: Electron density maps, GPC3-9-mer in TCR-unbound and TCR-bound states.**

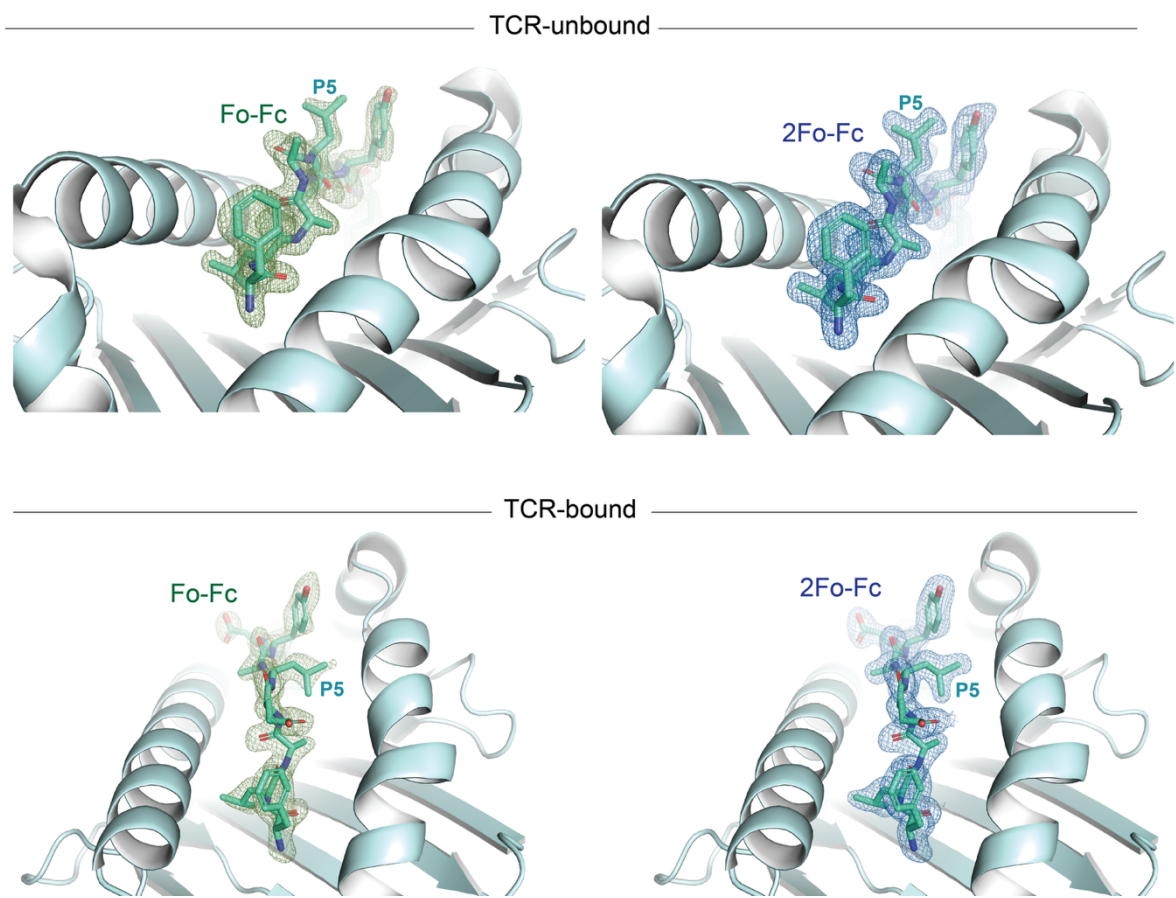

Fo-Fc and 2Fo-Fc electron density maps for the bound 9-mer peptide in its TCR-unbound and TCR-B-bound forms.

**Supplementary Figure 2: Electron density map of GPC3-11-mer, bound to HLA-A2.**

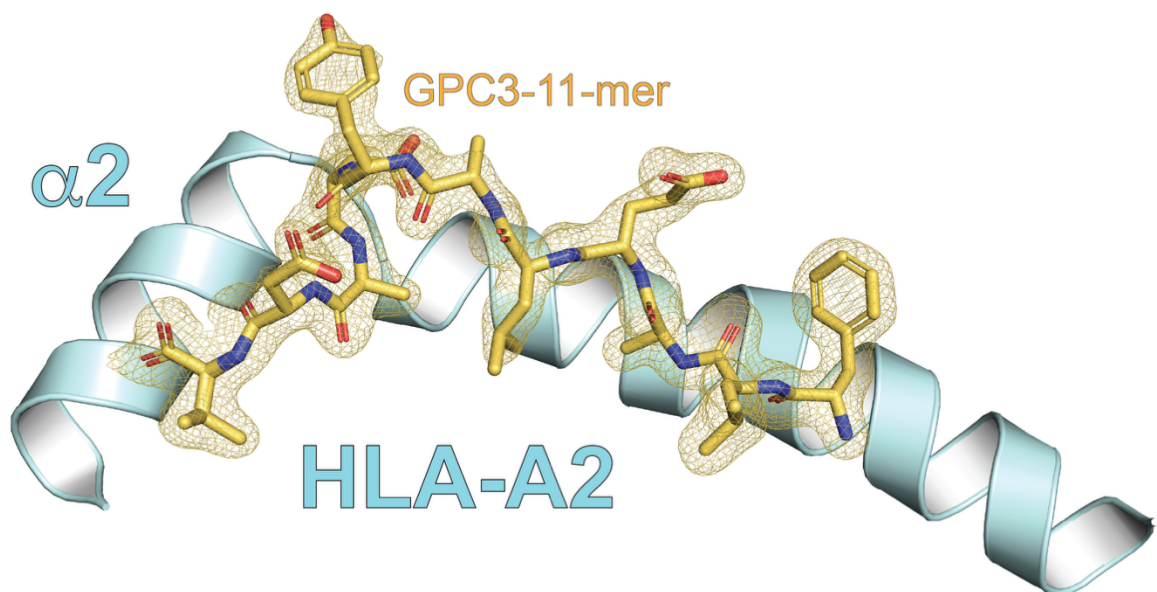

2Fo-Fc electron density map (contoured at 1 sigma) for the bound 11-mer peptide. For easy visualization, the MHC $\alpha 1$ and $\alpha 3$ domains are omitted.

**Supplementary Figure 3: Positional differences in the CDR3$\alpha/\beta$ loops near the peptide Y7.**

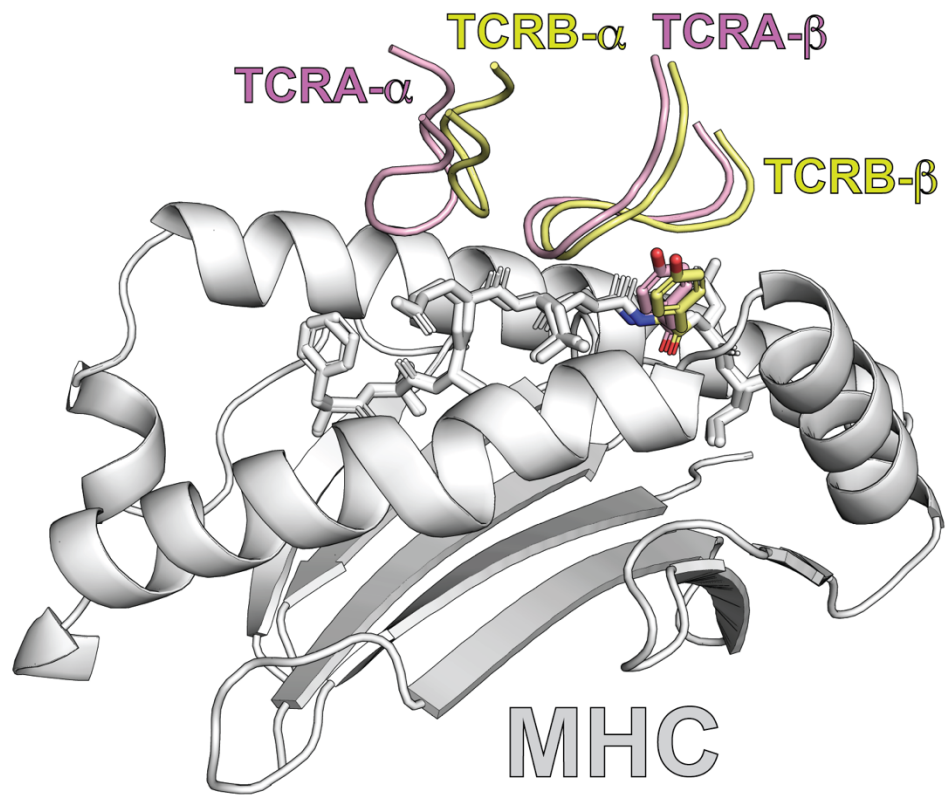

Structural alignment of the TCR-A (pink) and TCR-B (yellow) ternary complexes relative to the MHC $\alpha 1$-$\alpha 2$ platform (grey). CDR3α and CDR3β loops from each complex are shown near the peptide-binding interface, together with the peptide Y7 side chain (highlighted as sticks).

**Supplementary Figure 4: X-scan showing all tolerated ( $\geq 20\%$ ) changes for each FLAELAYDL position in TCR-A and TCR-B.**

Critical peptide motif for TCR-A reactivity

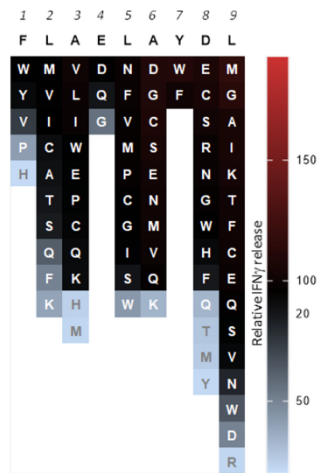

Critical peptide motif for TCR-B reactivity

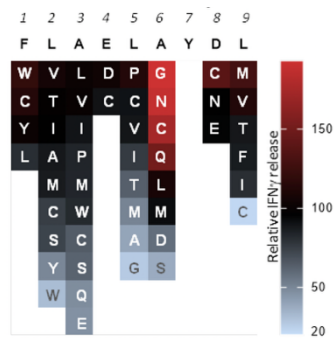

Amino acids are ranked in descending order based on their relative IFNγ release values.

**Supplementary Figure 5: Conjugate formation EO771 cell line.**

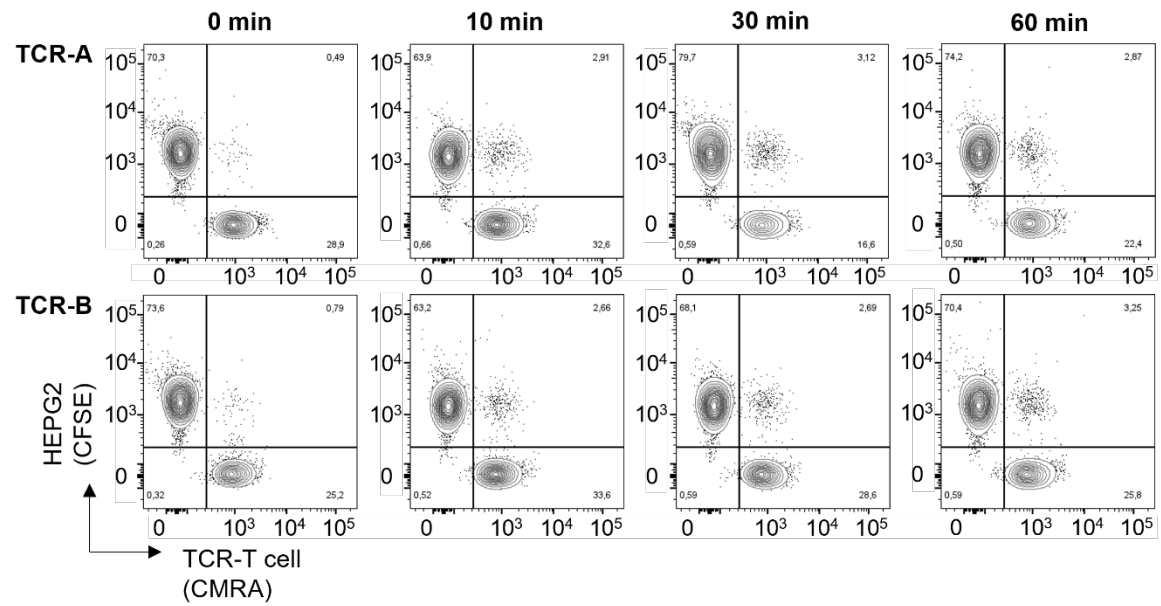

Representative flow cytometry plots showing conjugate formation between EO771 tumor cells and TCR-A or TCR-B T cells, identified as CFSE+ CMRA+ double-positive populations at the indicated time points.

**Supplementary Table 1: X-ray diffraction processing and refinement.**

|  | **HLA-A2:GPC3-9-mer** | **HLA-A2:GPC3-11-mer** | **TCR-A:HLA-A2:GPC3-9-mer** | **TCR-B:HLA-A2:GPC3-9-mer** |
| --- | --- | --- | --- | --- |
| Resolution range (Å) | 43.37 - 1.70 (1.761 - 1.7) | 32.26 - 1.8 (1.864 - 1.8) | 49.42 - 2.3 (2.382 - 2.3) | 30.3 - 1.95 (2.02 - 1.95) |
| Space group | P 1 21 1 | P 21 21 21 | P 65 2 2 | C 1 2 1 |
| Unit cell | 53.117 80.345 55.973 90 113.011 90 | 59.72 79.38 110.75 90 90 90 | 78.645 78.645 574.49 90 90 120 | 138.377 81.78 91.312 90 98.63 90 |
| Total reflections | 312057 (16076) | 324549 (31615) | 927459 (86743) | 239891 (14295) |
| Unique reflections | 47503 (2501) | 49524 (4891) | 48958 (4388) | 72498 (4463) |
| Multiplicity | 6.6 (6.4) | 6.6 (6.7) | 18.9 (19.8) | 3.3 (3.2) |
| Completeness (%) | 99.8 (99.8) | 100 (100) | 100.0 (100.00) | 99.0 (99.3) |
| Mean I/sigma(I) | 9.8 (1.1) | 11.7 (1.4) | 10.4 (2.7) | 6.2 (1.8) |
| Wilson B-factor | 23.43 | 29.77 | 36.22 | 22.67 |
| R-merge | 0.107 (1.776) | 0.075 (1.429) | 0.277 (1.568) | 0.156 (1.079) |
| R-meas | 0.116 (1.933) | 0.082 (1.549) | 0.284 (1.610) | 0.186 (1.3) |
| R-pim | 0.045 (0.755) | 0.032 (0.595) | 0.065 (0.360) | 0.101 (0.717) |
| CC1/2 | 0.998 (0.622) | 0.997 (0.600) | 0.997 (0.314) | 0.993 (0.302) |
| Reflections used in refinement | 47394 (4707) | 49517 (4891) | 48600 (4770) | 71652 (7131) |
| Reflections used for R-free | 2403 (228) | 2472 (248) | 2493 (236) | 3630 (346) |
| R-work | 0.1957 (0.3098) | 0.1702 (0.2193) | 0.2416 (0.3283) | 0.2597 (0.3640) |
| R-free | 0.2364 (0.3121) | 0.1983 (0.2637) | 0.2789 (0.3425) | 0.2898 (0.3812) |
| CC(work) | 0.961 (0.822) | 0.963 (0.914) | 0.928 (0.500) | 0.926 (0.381) |
| CC(free) | 0.952 (0.811) | 0.961 (0.869) | 0.905 (0.481) | 0.900 (0.378) |
| RMS(bonds) | 0.015 | 0.015 | 0.002 | 0.003 |
| RMS(angles) | 1.32 | 1.29 | 0.48 | 0.63 |
| Ramachandran favored (%) | 99.20 | 99.47 | 99.23 | 98.08 |
| Ramachandran allowed (%) | 0.80 | 0.53 | 0.77 | 1.92 |
| Ramachandran outliers (%) | 0.00 | 0.00 | 0.00 | 0.00 |
| Average B-factor (Å<sup>2</sup>) | 29.02 | 33.88 | 48.98 | 38.22 |

Statistics for the highest-resolution shell are shown in parentheses.

**Supplementary Table 2: HLA-A2 (chain A) with GPC3-9-mer (chain C) interactions found in 9TM0.** All intermolecular contacts are assigned according to the following distance cutoffs: hydrogen bonds $\leq 3.4$ Å, salt bridges $\leq 4.5$ Å, Van der Waals $\leq 4.0$ Å.

| HLA-A2 | GPC3-9-mer | Bond type |
| --- | --- | --- |
| Met5 S<sup>δ</sup> | Phe1 N | Van der Waals |
| Tyr7 C<sup>γ</sup> | Leu2 C<sup>δ2</sup> | Van der Waals |
| Tyr7 C<sup>δ1</sup> | Leu2 C<sup>δ2</sup> | Van der Waals |
| Tyr7 C<sup>δ2</sup> | Leu2 C<sup>δ2</sup> | Van der Waals |
| Tyr7 C<sup>ε1</sup> | Leu2 C<sup>δ2</sup> | Van der Waals |
| Tyr7 C<sup>ε2</sup> | Leu2 C<sup>δ2</sup>, Phe1 N, Phe1 O | Van der Waals |
| Tyr7 C<sup>ζ</sup> | Leu2 C<sup>δ2</sup>, Phe1 N | Van der Waals |
| Tyr7 O<sup>η</sup> | Phe1 N | Hydrogen bond |
| Tyr7 O<sup>η</sup> | Phe1 C<sup>α</sup>, Leu2 N, Phe1 C | Van der Waals |
| Phe9 C<sup>ζ</sup> | Leu2 C<sup>δ2</sup> | Van der Waals |
| Met45 C<sup>ε</sup> | Leu2 C<sup>δ1</sup> | Van der Waals |
| Glu63 O | Leu2 C<sup>δ1</sup> | Van der Waals |
| Glu63 C<sup>δ</sup> | Phe1 C<sup>α</sup>, Leu2 N | Van der Waals |
| Glu63 O<sup>ε1</sup> | Leu2 C<sup>β</sup>, Leu2 C<sup>γ</sup>, Leu2 C<sup>δ1</sup>, Phe1 C<sup>α</sup>, Leu2 C<sup>α</sup>, Phe1 C | Van der Waals |
| Glu63 O<sup>ε1</sup> | Leu2 N | Hydrogen bond |
| Glu63 O<sup>ε2</sup> | Phe1 C<sup>α</sup>, Leu2 N, Phe1 C<sup>ε1</sup>, Phe1 C<sup>γ</sup>, Phe1 C<sup>δ1</sup> | Van der Waals |
| Lys66 C<sup>β</sup> | Leu2 C<sup>β</sup>, Leu2 C<sup>δ1</sup> | Van der Waals |
| Lys66 C<sup>γ</sup> | Ala3 O | Van der Waals |
| Lys66 C<sup>δ</sup> | Leu2 O, Ala3 O, Glu4 C<sup>α</sup> | Van der Waals |
| Lys66 C<sup>ε</sup> | Leu2 C<sup>β</sup>, Leu2 O | Van der Waals |
| Lys66 N<sup>ζ</sup> | Leu2 N, Leu2 C, Phe1 C<sup>ε1</sup>, Phe1 C<sup>ζ</sup>, Phe1 C<sup>γ</sup>, Phe1 C<sup>δ1</sup>, Phe1 C<sup>δ2</sup>, Phe1 C<sup>ε2</sup> | Van der Waals |
| Lys66 N<sup>ζ</sup> | Leu2 O | Hydrogen bond |
| Val67 N | Leu2 C<sup>δ1</sup> | Van der Waals |
| Val67 C<sup>α</sup> | Leu2 C<sup>δ1</sup> | Van der Waals |
| Val67 C<sup>β</sup> | Leu2 C<sup>δ1</sup> | Van der Waals |
| His70 C<sup>γ</sup> | Ala6 C<sup>β</sup> | Van der Waals |
| His70 C<sup>δ2</sup> | Ala6 C<sup>β</sup> | Van der Waals |
| His70 C<sup>ε1</sup> | Ala3 O | Van der Waals |
| Thr73 C<sup>β</sup> | Ala6 O, Ala6 C<sup>β</sup> | Van der Waals |
| Thr73 O<sup>γ1</sup> | Ala6 C, Ala6 C<sup>β</sup> | Van der Waals |
| Thr73 O<sup>γ1</sup> | Ala6 O | Hydrogen bond |
| Thr73 C<sup>γ2</sup> | Tyr7 C, Asp8 N, Asp8 C<sup>α</sup>, Ala6 C, Ala6 O, Tyr7 O | Van der Waals |
| Asp77 C<sup>β</sup> | Leu9 C<sup>γ</sup>, Leu9 C<sup>δ2</sup> | Van der Waals |
| Asp77 C<sup>γ</sup> | Leu9 N, Leu9 C<sup>γ</sup>, Leu9 C<sup>δ2</sup> | Van der Waals |
| Asp77 O<sup>δ1</sup> | Asp8 C<sup>α</sup>, Asp8 C, Asp8 C<sup>β</sup>, Leu9 C<sup>γ</sup>, Leu9 C<sup>δ1</sup> | Van der Waals |
| Asp77 O<sup>δ1</sup> | Leu9 N | Hydrogen bond |
| Asp77 O<sup>δ2</sup> | Leu9 C<sup>γ</sup>, Leu9 C<sup>δ2</sup> | Van der Waals |
| Thr80 C<sup>β</sup> | Leu9 C<sup>δ1</sup> | Van der Waals |
| Thr80 C<sup>γ2</sup> | Leu9 O, Leu9 C<sup>δ1</sup> | Van der Waals |
| Leu81 C<sup>γ</sup> | Leu9 C<sup>δ1</sup> | Van der Waals |
| Leu81 C<sup>δ1</sup> | Leu9 C<sup>δ2</sup> | Van der Waals |
| Leu81 C<sup>δ2</sup> | Leu9 C<sup>δ1</sup> | Van der Waals |
| Tyr84 C<sup>ε2</sup> | Leu9 O | Van der Waals |
| Tyr84 C<sup>ζ</sup> | Leu9 O | Van der Waals |
| Tyr84 O<sup>η</sup> | Leu9 C, Leu9 O | Van der Waals |
| Tyr84 O<sup>η</sup> | Leu9 O | Hydrogen bond |
| Tyr99 C<sup>ζ</sup> | Ala3 N, Ala3 C<sup>β</sup> | Van der Waals |
| Tyr99 O<sup>η</sup> | Leu2 C<sup>β</sup>, Leu2 C<sup>δ2</sup>, Leu2 C<sup>α</sup>, Leu2 C, Ala3 C<sup>α</sup>, Ala3 C<sup>β</sup> | Van der Waals |
| Tyr99 O<sup>η</sup> | Ala3 N | Hydrogen bond |
| Tyr116 C<sup>ε1</sup> | Leu9 C<sup>δ2</sup> | Van der Waals |
| Tyr116 C<sup>ζ</sup> | Leu9 C<sup>δ2</sup> | Van der Waals |
| Tyr116 O<sup>η</sup> | Leu9 C<sup>δ2</sup> | Van der Waals |
| Tyr123 C<sup>ε2</sup> | Leu9 C<sup>δ2</sup> | Van der Waals |
| Thr143 C<sup>α</sup> | Leu9 O | Van der Waals |
| Thr143 C<sup>β</sup> | Leu9 O | Van der Waals |
| Thr143 O<sup>γ1</sup> | Leu9 C<sup>α</sup>, Leu9 C<sup>β</sup>, Leu9 C | Van der Waals |
| Thr143 O<sup>γ1</sup> | Leu9 O | Hydrogen bond |
| Thr143 C<sup>γ2</sup> | Leu9 C<sup>α</sup>, Leu9 C<sup>β</sup>, Leu9 O | Van der Waals |
| Lys146 C<sup>δ</sup> | Asp8 O | Van der Waals |
| Lys146 C<sup>ε</sup> | Leu9 O, Leu9 C | Van der Waals |
| Lys146 N<sup>ζ</sup> | Leu9 C, Leu9 O | Van der Waals |
| Lys146 N<sup>ζ</sup> | Asp8 O<sup>δ2</sup> | Salt bridge |
| Lys146 N<sup>ζ</sup> | Leu9 O | Hydrogen bond |
| Trp147 C<sup>δ1</sup> | Asp8 O | Van der Waals |
| Trp147 N<sup>ε1</sup> | Asp8 C, Leu9 C<sup>α</sup>, Tyr7 O, Tyr7 C<sup>β</sup> | Van der Waals |
| Trp147 N<sup>ε1</sup> | Asp8 O | Hydrogen bond |
| Trp147 C<sup>ε2</sup> | Tyr7 O, Asp8 O | Van der Waals |
| Trp147 C<sup>ζ2</sup> | Tyr7 O | Van der Waals |
| Val152 C<sup>γ1</sup> | Tyr7 C<sup>δ2</sup> | Van der Waals |
| Val152 C<sup>γ2</sup> | Tyr7 C<sup>β</sup>, Tyr7 C<sup>γ</sup>, Tyr7 C<sup>δ2</sup>, Tyr7 C<sup>ε2</sup> | Van der Waals |
| Gln155 C<sup>δ</sup> | Tyr7 O<sup>η</sup>, Tyr7 C<sup>ε2</sup> | Van der Waals |
| Gln155 O<sup>ε1</sup> | Leu5 C<sup>δ1</sup>, Tyr7 C<sup>ε2</sup>, Tyr7 C<sup>ζ</sup> | Van der Waals |
| Gln155 O<sup>ε1</sup> | Tyr7 O<sup>η</sup> | Hydrogen bond |
| Gln155 N<sup>ε2</sup> | Tyr7 O<sup>η</sup> | Hydrogen bond |
| Gln155 N<sup>ε2</sup> | Tyr7 C<sup>ε2</sup>, Tyr7 C<sup>ζ</sup> | Van der Waals |
| Tyr159 C<sup>δ1</sup> | Ala3 C<sup>α</sup> | Van der Waals |
| Tyr159 C<sup>ε1</sup> | Leu2 C, Leu2 O, Ala3 N, Ala3 C<sup>α</sup>, Phe1 O | Van der Waals |
| Tyr159 C<sup>ζ</sup> | Ala3 N, Ala3 C<sup>α</sup>, Phe1 O | Van der Waals |
| Tyr159 O<sup>η</sup> | Leu2 C<sup>α</sup>, Leu2 C, Ala3 N, Phe1 C | Van der Waals |
| Tyr159 O<sup>η</sup> | Phe1 O | Hydrogen bond |
| Thr163 C<sup>γ2</sup> | Phe1 C<sup>δ2</sup>, Phe1 C<sup>ε2</sup> | Van der Waals |
| Trp167 C<sup>β</sup> | Phe1 C<sup>β</sup> | Van der Waals |
| Trp167 C<sup>γ</sup> | Phe1 C<sup>β</sup>, Phe1 C<sup>γ</sup> | Van der Waals |
| Trp167 C<sup>δ1</sup> | Phe1 C<sup>β</sup>, Phe1 C<sup>γ</sup>, Phe1 C<sup>δ2</sup> | Van der Waals |
| Trp167 C<sup>δ2</sup> | Phe1 C<sup>β</sup>, Phe1 C<sup>γ</sup>, Phe1 C<sup>δ1</sup> | Van der Waals |
| Trp167 N<sup>ε1</sup> | Phe1 C<sup>ε1</sup>, Phe1 C<sup>ζ</sup>, Phe1 C<sup>γ</sup>, Phe1 C<sup>δ1</sup>, Phe1 C<sup>δ2</sup> | Van der Waals |
| Trp167 C<sup>ε2</sup> | Phe1 C<sup>ε1</sup>, Phe1 C<sup>γ</sup>, Phe1 C<sup>δ1</sup> | Van der Waals |
| Trp167 C<sup>ζ2</sup> | Phe1 C<sup>ε1</sup>, Phe1 C<sup>δ1</sup> | Van der Waals |
| Trp167 C<sup>η2</sup> | Phe1 C<sup>δ1</sup> | Van der Waals |
| Tyr171 C<sup>ε2</sup> | Phe1 N | Van der Waals |
| Tyr171 C<sup>ζ</sup> | Phe1 N | Van der Waals |
| Tyr171 O<sup>η</sup> | Phe1 N | Hydrogen bond |
| Tyr171 O<sup>η</sup> | Phe1 C<sup>α</sup>, Phe1 C<sup>β</sup> | Van der Waals |

**Supplementary Table 3: HLA-A2 (chain A) with GPC3-11-mer (chain P) interactions found in 31CS.** All intermolecular contacts are assigned according to the following distance cutoffs: hydrogen bonds $\leq 3.4$ Å, salt bridges $\leq 4.5$ Å, Van der Waals $\leq 4.0$ Å.

| HLA-A2 | GPC3-11-mer | Bond type |
| --- | --- | --- |
| Met5 S<sup>δ</sup> | Phe1 N | Van der Waals |
| Tyr7 C<sup>γ</sup> | Leu2 C<sup>δ2</sup> | Van der Waals |
| Tyr7 C<sup>δ1</sup> | Leu2 C<sup>δ2</sup> | Van der Waals |
| Tyr7 C<sup>δ2</sup> | Leu2 C<sup>δ2</sup> | Van der Waals |
| Tyr7 C<sup>ε1</sup> | Leu2 C<sup>γ</sup>, Leu2 C<sup>δ2</sup> | Van der Waals |
| Tyr7 C<sup>ε2</sup> | Phe1 N, Phe1 C, Phe1 O, Leu2 C<sup>δ2</sup> | Van der Waals |
| Tyr7 C<sup>ζ</sup> | Leu2 C<sup>γ</sup>, Phe1 N, Leu2 C<sup>δ2</sup> | Van der Waals |
| Tyr7 O<sup>η</sup> | Phe1 N | Hydrogen bond |
| Tyr7 O<sup>η</sup> | Phe1 C<sup>α</sup>, Phe1 C, Leu2 N | Van der Waals |
| Phe9 C<sup>ε2</sup> | Leu2 C<sup>δ2</sup> | Van der Waals |
| Phe9 C<sup>ζ</sup> | Leu2 C<sup>δ2</sup> | Van der Waals |
| Met45 C<sup>ε</sup> | Leu2 C<sup>γ</sup>, Leu2 C<sup>δ1</sup> | Van der Waals |
| Glu63 O | Leu2 C<sup>δ1</sup> | Van der Waals |
| Glu63 C<sup>δ</sup> | Phe1 C<sup>α</sup>, Leu2 N | Van der Waals |
| Glu63 O<sup>ε1</sup> | Leu2 C<sup>β</sup>, Leu2 C<sup>γ</sup>, Leu2 C<sup>δ1</sup>, Phe1 C<sup>α</sup>, Phe1 C, Leu2 C<sup>α</sup> | Van der Waals |
| Glu63 O<sup>ε1</sup> | Leu2 N | Hydrogen bond |
| Glu63 O<sup>ε2</sup> | Phe1 C<sup>γ</sup>, Phe1 C<sup>δ1</sup>, Phe1 C<sup>ε1</sup>, Phe1 C<sup>α</sup>, Leu2 N | Van der Waals |
| Arg65 C<sup>ζ</sup> | Glu4 O<sup>ε1</sup>, Glu4 O<sup>ε2</sup> | Van der Waals |
| Arg65 N<sup>η1</sup> | Glu4 O<sup>ε1</sup>, Glu4 O<sup>ε2</sup> | Salt bridge |
| Arg65 N<sup>η1</sup> | Glu4 C<sup>δ</sup> | Van der Waals |
| Arg65 N<sup>η2</sup> | Glu4 O<sup>ε1</sup>, Glu4 O<sup>ε2</sup> | Salt bridge |
| Arg65 N<sup>η2</sup> | Glu4 C<sup>δ</sup> | Van der Waals |
| Lys66 C<sup>β</sup> | Leu2 C<sup>β</sup>, Leu2 C<sup>δ1</sup> | Van der Waals |
| Lys66 C<sup>γ</sup> | Glu4 O<sup>ε1</sup>, Ala3 O, Glu4 C<sup>δ</sup>, Glu4 O<sup>ε2</sup> | Van der Waals |
| Lys66 C<sup>δ</sup> | Leu2 O, Ala3 O, Glu4 C<sup>δ</sup>, Glu4 O<sup>ε2</sup>, Ala3 C | Van der Waals |
| Lys66 C<sup>ε</sup> | Leu2 O, Leu2 C<sup>β</sup> | Van der Waals |
| Lys66 N<sup>ζ</sup> | Leu2 C, Phe1 C<sup>γ</sup>, Phe1 C<sup>δ1</sup>, Phe1 C<sup>δ2</sup>, Phe1 C<sup>ε1</sup>, Phe1 C<sup>ε2</sup>, Phe1 C<sup>ζ</sup>, Leu2 N | Van der Waals |
| Lys66 N<sup>ζ</sup> | Leu2 O | Hydrogen bond |
| Val67 N | Leu2 C<sup>δ1</sup> | Van der Waals |
| Val67 C<sup>α</sup> | Leu2 C<sup>δ1</sup> | Van der Waals |
| Val67 C<sup>β</sup> | Leu2 C<sup>δ1</sup> | Van der Waals |
| His70 C<sup>α</sup> | Leu5 C<sup>δ2</sup> | Van der Waals |
| His70 C<sup>β</sup> | Leu5 C<sup>δ2</sup> | Van der Waals |
| His70 C<sup>γ</sup> | Leu5 C<sup>δ2</sup> | Van der Waals |
| His70 N<sup>δ1</sup> | Leu5 C<sup>β</sup>, Leu5 C<sup>δ2</sup> | Van der Waals |
| His70 C<sup>δ2</sup> | Leu5 C<sup>δ2</sup> | Van der Waals |
| His70 C<sup>ε1</sup> | Ala3 O, Leu5 C<sup>β</sup> | Van der Waals |
| His70 N<sup>ε2</sup> | Ala3 O, Leu5 C<sup>δ2</sup> | Van der Waals |
| Thr73 C<sup>β</sup> | Asp10 O<sup>δ2</sup> | Van der Waals |
| Thr73 O<sup>γ1</sup> | Asp10 O<sup>δ2</sup>, Leu5 O | Van der Waals |
| Thr73 C<sup>γ2</sup> | Asp10 O<sup>δ2</sup>, Leu9 O | Van der Waals |
| Val76 C<sup>γ1</sup> | Asp10 O<sup>δ1</sup> | Van der Waals |
| Asp77 C<sup>β</sup> | Val11 C<sup>γ2</sup> | Van der Waals |
| Asp77 C<sup>γ</sup> | Val11 N, Val11 C<sup>γ1</sup>, Val11 C<sup>γ2</sup> | Van der Waals |
| Asp77 O<sup>δ1</sup> | Asp10 C<sup>α</sup>, Asp10 C, Asp10 C<sup>β</sup>, Val11 C<sup>α</sup>, Val11 C<sup>β</sup>, Val11 C<sup>γ1</sup>, Val11 C<sup>γ2</sup> | Van der Waals |
| Asp77 O<sup>δ1</sup> | Val11 N | Hydrogen bond |
| Asp77 O<sup>δ2</sup> | Val11 C<sup>γ1</sup> | Van der Waals |
| Thr80 C<sup>γ2</sup> | Val11 O | Van der Waals |
| Leu81 C<sup>γ</sup> | Val11 C<sup>γ2</sup> | Van der Waals |
| Tyr84 C<sup>ε2</sup> | Val11 O | Van der Waals |
| Tyr84 C<sup>5</sup> | Val11 O | Van der Waals |
| Tyr84 O<sup>n</sup> | Val11 C, Val11 O | Van der Waals |
| Tyr84 O<sup>n</sup> | Val11 O | Hydrogen bond |
| Arg97 C<sup>5</sup> | Leu5 C<sup>δ2</sup> | Van der Waals |
| Tyr99 C<sup>5</sup> | Ala3 N, Ala3 C<sup>β</sup> | Van der Waals |
| Tyr99 O<sup>n</sup> | Leu2 C, Leu2 C<sup>β</sup>, Leu2 C<sup>α</sup>, Ala3 C<sup>α</sup>, Ala3 C<sup>β</sup>,<br>Leu2 C<sup>δ2</sup> | Van der Waals |
| Tyr99 O<sup>n</sup> | Ala3 N | Hydrogen bond |
| Tyr116 C<sup>ε2</sup> | Val11 C<sup>γ1</sup> | Van der Waals |
| Thr143 C<sup>α</sup> | Val11 O | Van der Waals |
| Thr143 C<sup>β</sup> | Val11 O | Van der Waals |
| Thr143 O<sup>γ1</sup> | Val11 C<sup>α</sup>, Val11 C, Val11 C<sup>β</sup> | Van der Waals |
| Thr143 O<sup>γ1</sup> | Val11 O | Hydrogen bond |
| Thr143 C<sup>γ2</sup> | Val11 C<sup>α</sup>, Val11 C<sup>β</sup>, Val11 C<sup>γ1</sup>, Val11 O | Van der Waals |
| Trp147 C<sup>δ1</sup> | Asp8 O, Asp10 O | Van der Waals |
| Trp147 N<sup>ε1</sup> | Asp8 O, Asp10 C, Leu9 C | Van der Waals |
| Trp147 N<sup>ε1</sup> | Asp10 O | Hydrogen bond |
| Trp147 C<sup>ε2</sup> | Asp10 O, Leu9 O | Van der Waals |
| Trp147 C<sup>ζ2</sup> | Val11 C<sup>γ1</sup>, Leu9 O | Van der Waals |
| Val152 C<sup>γ2</sup> | Asp8 O | Van der Waals |
| Gln155 C<sup>δ</sup> | Asp8 O<sup>δ1</sup> | Van der Waals |
| Gln155 O<sup>ε1</sup> | Asp8 O<sup>δ1</sup> | Van der Waals |
| Gln155 N<sup>ε2</sup> | Asp8 C<sup>γ</sup>, Asp8 O<sup>δ1</sup> | Van der Waals |
| Tyr159 C<sup>γ</sup> | Ala3 C<sup>β</sup> | Van der Waals |
| Tyr159 C<sup>δ1</sup> | Ala3 C<sup>α</sup>, Ala3 C<sup>β</sup> | Van der Waals |
| Tyr159 C<sup>δ2</sup> | Ala3 C<sup>β</sup> | Van der Waals |
| Tyr159 C<sup>ε1</sup> | Leu2 C, Leu2 O, Phe1 O, Ala3 N, Ala3 C<sup>α</sup>,<br>Ala3 C<sup>β</sup> | Van der Waals |
| Tyr159 C<sup>ε2</sup> | Ala3 C<sup>β</sup> | Van der Waals |
| Tyr159 C<sup>5</sup> | Phe1 O, Ala3 N, Ala3 C<sup>α</sup>, Ala3 C<sup>β</sup> | Van der Waals |
| Tyr159 O<sup>n</sup> | Leu2 C, Phe1 C, Ala3 N | Van der Waals |
| Tyr159 O<sup>n</sup> | Phe1 O | Hydrogen bond |
| Thr163 C<sup>γ2</sup> | Phe1 C<sup>δ2</sup>, Phe1 C<sup>ε2</sup> | Van der Waals |
| Trp167 C<sup>β</sup> | Phe1 C<sup>β</sup> | Van der Waals |
| Trp167 C<sup>γ</sup> | Phe1 C<sup>β</sup>, Phe1 C<sup>γ</sup> | Van der Waals |
| Trp167 C<sup>δ1</sup> | Phe1 C<sup>β</sup>, Phe1 C<sup>γ</sup>, Phe1 C<sup>δ2</sup> | Van der Waals |
| Trp167 C<sup>δ2</sup> | Phe1 C<sup>β</sup>, Phe1 C<sup>γ</sup>, Phe1 C<sup>δ1</sup> | Van der Waals |
| Trp167 N<sup>ε1</sup> | Phe1 C<sup>γ</sup>, Phe1 C<sup>δ1</sup>, Phe1 C<sup>δ2</sup>, Phe1 C<sup>ε1</sup>, Phe1 C<sup>ε2</sup> | Van der Waals |
| Trp167 C<sup>ε2</sup> | Phe1 C<sup>γ</sup>, Phe1 C<sup>δ1</sup>, Phe1 C<sup>ε1</sup> | Van der Waals |
| Trp167 C<sup>ζ2</sup> | Phe1 C<sup>δ1</sup>, Phe1 C<sup>ε1</sup> | Van der Waals |
| Trp167 C<sup>η2</sup> | Phe1 C<sup>δ1</sup> | Van der Waals |
| Tyr171 C<sup>ε2</sup> | Phe1 N | Van der Waals |
| Tyr171 C<sup>5</sup> | Phe1 N | Van der Waals |
| Tyr171 O<sup>n</sup> | Phe1 C<sup>β</sup>, Phe1 C<sup>α</sup> | Van der Waals |
| Tyr171 O<sup>n</sup> | Phe1 N | Hydrogen bond |

**Supplementary Table 4: TCR-A $\alpha$ (chain C) with HLA-A2 (chain A) interactions found in 9TMM.** All intermolecular contacts are assigned according to the following distance cutoffs: hydrogen bonds $\leq 3.4$ Å, salt bridges $\leq 4.5$ Å, Van der Waals $\leq 4.0$ Å.

| TCR-A $\alpha$ | HLA-A2 | Bond type |
| --- | --- | --- |
| Ser28 C | Trp167 N<sup>e1</sup> | Van der Waals |
| Ser28 O | Trp167 C<sup>δ1</sup>, Trp167 C<sup>ε2</sup> | Van der Waals |
| Ser28 O | Trp167 N<sup>e1</sup> | Hydrogen bond |
| Ser28 O<sup>γ</sup> | Trp167 C<sup>ε2</sup> | Van der Waals |
| Gly29 C<sup>α</sup> | Thr163 C<sup>γ2</sup> | Van der Waals |
| Gly29 C | Thr163 C<sup>γ2</sup>, Thr163 O<sup>γ1</sup> | Van der Waals |
| Thr30 N | Thr163 C<sup>γ2</sup> | Van der Waals |
| Thr30 N | Thr163 O<sup>γ1</sup> | Hydrogen bond |
| Thr30 C<sup>α</sup> | Thr163 O<sup>γ1</sup> | Van der Waals |
| Thr30 C<sup>β</sup> | Thr163 O<sup>γ1</sup> | Van der Waals |
| Thr30 O<sup>γ1</sup> | Tyr159 C<sup>β</sup>, Tyr159 C<sup>α</sup>, Tyr159 C<sup>δ1</sup>, Thr163 C<sup>β</sup> | Van der Waals |
| Thr30 O<sup>γ1</sup> | Thr163 O<sup>γ1</sup> | Hydrogen bond |
| Thr30 C<sup>γ2</sup> | Thr163 O<sup>γ1</sup>, Ala158 C, Ala158 O, Ala158 C<sup>β</sup>, Tyr159 N | Van der Waals |
| Tyr32 C<sup>ε1</sup> | Gln155 O | Van der Waals |
| Tyr32 C<sup>ζ</sup> | Gln155 O | Van der Waals |
| Tyr32 O<sup>η</sup> | Gln155 C | Van der Waals |
| Tyr32 O<sup>η</sup> | Gln155 O | Hydrogen bond |
| Tyr51 O | Ala158 C<sup>β</sup> | Van der Waals |
| Tyr51 C<sup>δ1</sup> | Glu154 O, Gln155 C<sup>α</sup> | Van der Waals |
| Tyr51 C<sup>ε1</sup> | Glu154 C, Glu154 O, Gln155 N, Gln155 C<sup>α</sup>, Glu154 C<sup>β</sup>, Gln155 C<sup>γ</sup> | Van der Waals |
| Tyr51 C<sup>ε2</sup> | Gln155 C<sup>γ</sup> | Van der Waals |
| Tyr51 C<sup>ζ</sup> | Gln155 C<sup>γ</sup> | Van der Waals |
| Tyr51 O<sup>η</sup> | His151 C<sup>β</sup>, Gln155 C<sup>γ</sup> | Van der Waals |
| Ser52 C<sup>β</sup> | Ala158 C<sup>β</sup> | Van der Waals |
| Ser52 O<sup>γ</sup> | Glu154 O | Van der Waals |
| Lys69 C<sup>ε</sup> | Glu166 O<sup>ε2</sup> | Van der Waals |
| Lys69 N<sup>ζ</sup> | Glu166 O<sup>ε2</sup> | Salt bridge |
| Asn95 O | Arg65 N<sup>η2</sup> | Van der Waals |
| Asn96 C | Arg65 N<sup>ε</sup> | Van der Waals |
| Asn96 O | Arg65 C<sup>γ</sup>, Arg65 C<sup>δ</sup>, Arg65 C<sup>ζ</sup>, Arg65 N<sup>η2</sup> | Van der Waals |
| Asn96 O | Arg65 N<sup>ε</sup> | Hydrogen bond |
| Asn96 C<sup>β</sup> | Arg65 N<sup>η2</sup>, Gly62 C<sup>α</sup> | Van der Waals |
| Asn96 C<sup>γ</sup> | Gly62 C<sup>α</sup> | Van der Waals |
| Asn96 O<sup>δ1</sup> | Lys66 C<sup>ε</sup>, Lys66 C<sup>δ</sup> | Van der Waals |
| Asn96 O<sup>δ1</sup> | Lys66 N<sup>ζ</sup> | Hydrogen bond |
| Asn96 N<sup>δ2</sup> | Gly62 C<sup>α</sup> | Van der Waals |
| Asn97 O<sup>δ1</sup> | Lys66 C<sup>γ</sup> | Van der Waals |
| Asn97 N<sup>δ2</sup> | Ala69 C<sup>β</sup> | Van der Waals |
| Ala98 N | Arg65 N<sup>ε</sup> | Van der Waals |
| Ala98 C<sup>β</sup> | Arg65 N<sup>ε</sup>, Arg65 C<sup>ζ</sup>, Arg65 N<sup>η2</sup> | Van der Waals |

**Supplementary Table 5: TCR-A $\beta$ (chain D) with HLA-A2 (chain A) interactions found in 9TMM.** All intermolecular contacts are assigned according to the following distance cutoffs: hydrogen bonds $\leq 3.4$ Å, salt bridges $\leq 4.5$ Å, Van der Waals $\leq 4.0$ Å.

| TCR-A $\beta$ | HLA-A2 | Bond type |
| --- | --- | --- |
| Trp32 C$^{\zeta 2}$ | Thr73 O$^{\gamma 1}$ | Van der Waals |
| Trp32 C$^{\eta 2}$ | Thr73 O$^{\gamma 1}$ | Van der Waals |
| Arg51 N | Gln72 N$^{\epsilon 2}$ | Van der Waals |
| Arg51 C$^{\beta}$ | Gln72 N$^{\epsilon 2}$ | Van der Waals |
| Arg51 C$^{\gamma}$ | Val76 C$^{\gamma 2}$ | Van der Waals |
| Arg51 N$^{\eta 2}$ | Val76 C$^{\gamma 1}$ | Van der Waals |
| Ser52 O$^{\gamma}$ | Val76 C$^{\gamma 2}$ | Van der Waals |
| Asp55 O$^{\delta 2}$ | Gln72 N$^{\epsilon 2}$ | Hydrogen bond |
| Gly99 C$^{\alpha}$ | Gln155 O$^{\epsilon 1}$ | Van der Waals |
| Gly99 C | Gln155 O$^{\epsilon 1}$ | Van der Waals |
| Gly100 N | Gln155 O$^{\epsilon 1}$ | Hydrogen bond |
| Gly100 N | Gln155 C$^{\delta}$ | Van der Waals |
| Gly100 C$^{\alpha}$ | Gln155 O$^{\epsilon 1}$ | Van der Waals |
| Gly100 C | Gln155 O$^{\epsilon 1}$ | Van der Waals |
| Ala101 N | Gln155 O$^{\epsilon 1}$ | Van der Waals |

**Supplementary Table 6: TCR-A $\alpha$ (chain C) with GPC3-9-mer (chain P) interactions found in 9TMM.** All intermolecular contacts are assigned according to the following distance cutoffs: hydrogen bonds $\leq 3.4$ Å, salt bridges $\leq 4.5$ Å, Van der Waals $\leq 4.0$ Å.

| TCR-A $\alpha$ | GPC3-9-mer | Bond type |
| --- | --- | --- |
| Tyr27 C<sup>ε2</sup> | Glu4 O<sup>ε2</sup> | Van der Waals |
| Tyr27 C<sup>ζ</sup> | Glu4 O<sup>ε2</sup> | Van der Waals |
| Tyr27 O<sup>η</sup> | Phe1 C<sup>ζ</sup>, Glu4 C<sup>γ</sup>, Glu4 C<sup>δ</sup> | Van der Waals |
| Tyr27 O<sup>η</sup> | Glu4 O<sup>ε2</sup> | Hydrogen bond |
| Gly29 C<sup>α</sup> | Glu4 O<sup>ε2</sup> | Van der Waals |
| Gly29 C | Glu4 O<sup>ε2</sup> | Van der Waals |
| Thr30 N | Glu4 C<sup>δ</sup> | Van der Waals |
| Thr30 N | Glu4 O<sup>ε2</sup> | Hydrogen bond |
| Thr30 C<sup>α</sup> | Glu4 O<sup>ε2</sup> | Van der Waals |
| Thr30 O | Glu4 O<sup>ε2</sup> | Van der Waals |
| Thr30 C<sup>β</sup> | Glu4 C<sup>δ</sup>, Glu4 O<sup>ε1</sup>, Glu4 O<sup>ε2</sup> | Van der Waals |
| Thr30 O<sup>γ1</sup> | Glu4 C<sup>δ</sup> | Van der Waals |
| Thr30 O<sup>γ1</sup> | Glu4 O<sup>ε1</sup>, Glu4 O<sup>ε2</sup> | Hydrogen bond |
| Tyr32 C<sup>ε2</sup> | Glu4 O, Glu4 O<sup>ε1</sup> | Van der Waals |
| Asn96 O<sup>δ1</sup> | Phe1 C<sup>ε1</sup>, Phe1 C<sup>ε2</sup>, Phe1 C<sup>ζ</sup>, Glu4 C<sup>γ</sup> | Van der Waals |
| Asn97 O<sup>δ1</sup> | Glu4 C<sup>α</sup>, Glu4 C, Glu4 O, Glu4 C<sup>β</sup> | Van der Waals |
| Asn97 O<sup>δ1</sup> | Leu5 N | Hydrogen bond |

**Supplementary Table 7: TCR-A $\beta$ (chain D) with GPC3-9-mer (chain P) interactions found in 9TMM.** All intermolecular contacts are assigned according to the following distance cutoffs: hydrogen bonds $\leq 3.4$ Å, salt bridges $\leq 4.5$ Å, Van der Waals $\leq 4.0$ Å.

| TCR-A $\beta$ | GPC3-9-mer | Bond type |
| --- | --- | --- |
| Trp32 C$^{\delta 1}$ | Asp8 O$^{\delta 1}$ | Van der Waals |
| Trp32 N$^{\epsilon 1}$ | Asp8 C$^{\gamma}$, Asp8 O$^{\delta 2}$ | Van der Waals |
| Trp32 N$^{\epsilon 1}$ | Asp8 O$^{\delta 1}$ | Hydrogen bond |
| Trp32 C$^{\epsilon 2}$ | Asp8 O$^{\delta 1}$ | Van der Waals |
| Arg51 C$^{\delta}$ | Asp8 O$^{\delta 2}$ | Van der Waals |
| Arg51 N$^{\epsilon}$ | Asp8 C$^{\gamma}$ | Van der Waals |
| Arg51 N$^{\epsilon}$ | Asp8 O$^{\delta 2}$ | Salt bridge |
| Arg51 C$^{\zeta}$ | Asp8 O$^{\delta 2}$ | Van der Waals |
| Arg51 N$^{\eta 2}$ | Asp8 O$^{\delta 2}$ | Salt bridge |
| Ala95 C | Tyr7 C$^{\epsilon 2}$ | Van der Waals |
| Ala95 O | Asp8 O$^{\delta 1}$, Tyr7 C$^{\delta 2}$ | Van der Waals |
| Gly96 N | Tyr7 C$^{\epsilon 2}$ | Van der Waals |
| Gly96 C$^{\alpha}$ | Ala6 O, Tyr7 C$^{\delta 2}$, Tyr7 C$^{\epsilon 2}$ | Van der Waals |
| Gly96 C | Ala6 O | Van der Waals |
| Thr97 N | Ala6 O | Hydrogen bond |
| Thr97 O | Ala6 N | Hydrogen bond |
| Thr97 O | Ala6 C$^{\alpha}$, Ala6 O, Leu5 C$^{\alpha}$, Leu5 C, Leu5 C$^{\beta}$ | Van der Waals |
| Thr97 C$^{\gamma 2}$ | Ala6 C$^{\beta}$, Ala6 O | Van der Waals |
| Gly98 C$^{\alpha}$ | Glu4 O | Van der Waals |
| Gly98 C | Glu4 O | Van der Waals |
| Gly99 N | Glu4 C, Leu5 C$^{\alpha}$, Leu5 C$^{\beta}$ | Van der Waals |
| Gly99 N | Glu4 O | Hydrogen bond |
| Gly99 C$^{\alpha}$ | Glu4 O, Leu5 C$^{\beta}$ | Van der Waals |
| Gly100 C | Tyr7 O$^{\eta}$ | Van der Waals |
| Ala101 N | Tyr7 O$^{\eta}$ | Hydrogen bond |
| Ala101 C$^{\alpha}$ | Tyr7 O$^{\eta}$ | Van der Waals |
| Ala101 C$^{\beta}$ | Tyr7 O$^{\eta}$ | Van der Waals |

**Supplementary Table 8: HLA-A2 (chain A) with GPC3-9-mer (chain P) interactions found in 9TMM.** All intermolecular contacts are assigned according to the following distance cutoffs: hydrogen bonds $\leq 3.4$ Å, salt bridges $\leq 4.5$ Å, Van der Waals $\leq 4.0$ Å.

| HLA-A2 | GPC3-9-mer | Bond type |
| --- | --- | --- |
| Tyr7 Cγ | Leu2 C$\delta^2$ | Van der Waals |
| Tyr7 C$\delta^1$ | Leu2 C$\delta^2$ | Van der Waals |
| Tyr7 C$\epsilon^1$ | Leu2 C$\delta^2$ | Van der Waals |
| Tyr7 C$\epsilon^2$ | Phe1 N, Phe1 C, Phe1 O | Van der Waals |
| Tyr7 Cζ | Leu2 C$\delta^2$, Phe1 N | Van der Waals |
| Tyr7 Oη | Leu2 N, Phe1 Cα, Phe1 C | Van der Waals |
| Tyr7 Oη | Phe1 N | Hydrogen bond |
| Phe9 Cζ | Leu2 C$\delta^2$ | Van der Waals |
| Met45 Cε | Leu2 C$\delta^1$ | Van der Waals |
| Glu63 O | Leu2 C$\delta^1$ | Van der Waals |
| Glu63 Cδ | Leu2 N, Phe1 Cα | Van der Waals |
| Glu63 O$\epsilon^1$ | Leu2 N | Hydrogen bond |
| Glu63 O$\epsilon^1$ | Leu2 Cα, Leu2 Cβ, Leu2 Cγ, Phe1 Cα, Phe1 C | Van der Waals |
| Glu63 O$\epsilon^2$ | Leu2 N, Phe1 Cγ, Phe1 C$\delta^1$, Phe1 C$\epsilon^1$, Phe1 Cα | Van der Waals |
| Lys66 C | Leu2 C$\delta^1$ | Van der Waals |
| Lys66 Cβ | Leu2 C$\delta^1$ | Van der Waals |
| Lys66 Cδ | Leu2 O, Glu4 Cα, Glu4 Cβ, Glu4 Cγ | Van der Waals |
| Lys66 Cε | Leu2 Cβ, Leu2 O | Van der Waals |
| Lys66 Nζ | Leu2 N, Leu2 O | Hydrogen bond |
| Lys66 Nζ | Leu2 Cα, Leu2 C, Leu2 Cβ, Phe1 Cγ, Phe1 C$\delta^1$, Phe1 C$\delta^2$, Phe1 C$\epsilon^1$, Phe1 C$\epsilon^2$, Phe1 Cζ | Van der Waals |
| Val67 N | Leu2 C$\delta^1$ | Van der Waals |
| Val67 Cα | Leu2 C$\delta^1$ | Van der Waals |
| Val67 Cβ | Leu2 C$\delta^1$ | Van der Waals |
| His70 C$\epsilon^1$ | Leu5 O, Ala3 O | Van der Waals |
| Thr73 O$\gamma^1$ | Ala6 Cβ | Van der Waals |
| Thr73 C$\gamma^2$ | Tyr7 C, Asp8 N, Asp8 Cα, Tyr7 O | Van der Waals |
| Val76 C$\gamma^1$ | Asp8 O$\delta^2$ | Van der Waals |
| Asp77 Cβ | Leu9 C$\delta^1$ | Van der Waals |
| Asp77 Cγ | Leu9 Cβ, Leu9 C$\delta^1$, Leu9 N, Leu9 Cγ | Van der Waals |
| Asp77 O$\delta^1$ | Leu9 Cβ, Asp8 Cα, Leu9 Cα, Leu9 Cγ, Asp8 C | Van der Waals |
| Asp77 O$\delta^1$ | Leu9 N | Hydrogen bond |
| Asp77 O$\delta^2$ | Leu9 C$\delta^1$, Leu9 Cγ | Van der Waals |
| Thr80 C$\gamma^2$ | Leu9 O | Van der Waals |
| Leu81 C$\delta^1$ | Leu9 C$\delta^1$ | Van der Waals |
| Tyr84 Oη | Leu9 O | Van der Waals |
| Tyr99 Cζ | Ala3 N, Ala3 Cβ | Van der Waals |
| Tyr99 Oη | Leu2 Cα, Leu2 C, Leu2 Cβ, Leu2 C$\delta^2$, Ala3 Cα, Ala3 Cβ | Van der Waals |
| Tyr99 Oη | Ala3 N | Hydrogen bond |
| Tyr116 Cγ | Leu9 C$\delta^2$ | Van der Waals |
| Tyr116 C$\delta^2$ | Leu9 C$\delta^1$, Leu9 C$\delta^2$ | Van der Waals |
| Tyr116 C$\epsilon^2$ | Leu9 C$\delta^2$ | Van der Waals |
| Tyr116 Cζ | Leu9 C$\delta^2$ | Van der Waals |
| Thr143 O$\gamma^1$ | Leu9 Cβ, Leu9 C, Leu9 Cα | Van der Waals |
| Thr143 C$\gamma^2$ | Leu9 C, Leu9 Cα, Leu9 C$\delta^2$ | Van der Waals |
| Lys146 Cε | Leu9 C, Leu9 O | Van der Waals |
| Lys146 Nζ | Leu9 C, Asp8 O | Van der Waals |
| Lys146 Nζ | Leu9 O | Hydrogen bond |
| Trp147 C$\delta^1$ | Asp8 O | Van der Waals |
| Trp147 N$\epsilon^1$ | Asp8 C | Van der Waals |
| Trp147 N$\epsilon^1$ | Asp8 O | Hydrogen bond |
| Trp147 C$\epsilon^2$ | Asp8 O | Van der Waals |
| Trp147 C<sup>ε2</sup> | Tyr7 O, Leu9 C<sup>α</sup>, Leu9 C<sup>δ2</sup>, Asp8 O | Van der Waals |
| Trp147 C<sup>η2</sup> | Leu9 C<sup>δ2</sup> | Van der Waals |
| Val152 C<sup>γ2</sup> | Tyr7 C<sup>γ</sup>, Tyr7 C<sup>δ1</sup> | Van der Waals |
| Gln155 C<sup>β</sup> | Leu5 C<sup>δ1</sup> | Van der Waals |
| Gln155 C<sup>δ</sup> | Tyr7 C<sup>ε1</sup>, Tyr7 O<sup>η</sup> | Van der Waals |
| Gln155 O<sup>ε1</sup> | Tyr7 C<sup>ε1</sup>, Tyr7 C<sup>ζ</sup> | Van der Waals |
| Gln155 O<sup>ε1</sup> | Tyr7 O<sup>η</sup> | Hydrogen bond |
| Gln155 N<sup>ε2</sup> | Tyr7 C<sup>ε1</sup>, Tyr7 C<sup>ζ</sup> | Van der Waals |
| Gln155 N<sup>ε2</sup> | Tyr7 O<sup>η</sup> | Hydrogen bond |
| Leu156 C<sup>δ2</sup> | Leu5 C<sup>δ2</sup> | Van der Waals |
| Tyr159 C<sup>γ</sup> | Ala3 C<sup>β</sup> | Van der Waals |
| Tyr159 C<sup>δ1</sup> | Ala3 C<sup>α</sup>, Ala3 C<sup>β</sup>, Glu4 O<sup>ε1</sup> | Van der Waals |
| Tyr159 C<sup>δ2</sup> | Ala3 C<sup>β</sup> | Van der Waals |
| Tyr159 C<sup>ε1</sup> | Leu2 C, Leu2 O, Ala3 N, Ala3 C<sup>α</sup>, Ala3 C<sup>β</sup>, Phe1 O, | Van der Waals |
| Tyr159 C<sup>ε2</sup> | Ala3 C<sup>β</sup> | Van der Waals |
| Tyr159 C<sup>ζ</sup> | Ala3 N, Ala3 C<sup>α</sup>, Ala3 C<sup>β</sup>, Phe1 O | Van der Waals |
| Tyr159 O<sup>η</sup> | Ala3 N, Phe1 C | Van der Waals |
| Tyr159 O<sup>η</sup> | Phe1 O | Hydrogen bond |
| Thr163 C<sup>γ2</sup> | Phe1 C<sup>δ2</sup>, Phe1 C<sup>ε2</sup> | Van der Waals |
| Trp167 C<sup>β</sup> | Phe1 C<sup>β</sup> | Van der Waals |
| Trp167 C<sup>γ</sup> | Phe1 C<sup>γ</sup>, Phe1 C<sup>β</sup> | Van der Waals |
| Trp167 C<sup>δ1</sup> | Phe1 C<sup>γ</sup>, Phe1 C<sup>β</sup> | Van der Waals |
| Trp167 C<sup>δ2</sup> | Phe1 C<sup>γ</sup>, Phe1 C<sup>δ1</sup>, Phe1 C<sup>β</sup> | Van der Waals |
| Trp167 N<sup>ε1</sup> | Phe1 C<sup>γ</sup>, Phe1 C<sup>δ1</sup> | Van der Waals |
| Trp167 C<sup>ε2</sup> | Phe1 C<sup>γ</sup>, Phe1 C<sup>δ1</sup>, Phe1 C<sup>ε1</sup> | Van der Waals |
| Trp167 C<sup>ε3</sup> | Phe1 C<sup>δ1</sup>, Phe1 C<sup>β</sup> | Van der Waals |
| Trp167 C<sup>ζ2</sup> | Phe1 C<sup>δ1</sup>, Phe1 C<sup>ε1</sup> | Van der Waals |
| Trp167 C<sup>ζ3</sup> | Phe1 C<sup>δ1</sup> | Van der Waals |
| Trp167 C<sup>η2</sup> | Phe1 C<sup>δ1</sup> | Van der Waals |
| Tyr171 C<sup>ε2</sup> | Phe1 N | Van der Waals |
| Tyr171 C<sup>ζ</sup> | Phe1 N | Van der Waals |
| Tyr171 O<sup>η</sup> | Phe1 N | Hydrogen bond |
| Tyr171 O<sup>η</sup> | Phe1 C<sup>α</sup>, Phe1 C<sup>β</sup> | Van der Waals |

**Supplementary Table 9: TCR-B $\alpha$ (chain C) with HLA-A2 (chain A) interactions found in 9TML.** All intermolecular contacts are assigned according to the following distance cutoffs: hydrogen bonds $\leq 3.4$ Å, salt bridges $\leq 4.5$ Å, Van der Waals $\leq 4.0$ Å.

| TCR-B $\alpha$ | HLA-A2 | Bond type |
| --- | --- | --- |
| Ser28 O | Trp167 C<sup>ε2</sup>, Trp167 C<sup>ζ2</sup> | Van der Waals |
| Ser28 O | Trp167 N<sup>ε1</sup> | Hydrogen bond |
| Ser28 C<sup>β</sup> | Trp167 C<sup>ζ2</sup> | Van der Waals |
| Ser28 O<sup>γ</sup> | Trp167 C<sup>ζ2</sup> | Van der Waals |
| Gly29 C<sup>α</sup> | Thr163 C<sup>γ2</sup> | Van der Waals |
| Gly29 C | Thr163 O<sup>γ1</sup>, Thr163 C<sup>γ2</sup> | Van der Waals |
| Gly29 O | Thr163 O<sup>γ1</sup> | Van der Waals |
| Thr30 N | Thr163 O<sup>γ1</sup> | Hydrogen bond |
| Thr30 C<sup>α</sup> | Thr163 O<sup>γ1</sup> | Van der Waals |
| Thr30 C<sup>β</sup> | Thr163 O<sup>γ1</sup> | Van der Waals |
| Thr30 O<sup>γ1</sup> | Thr163 C<sup>β</sup>, Thr163 C<sup>γ2</sup>, Tyr159 C<sup>β</sup>, Tyr159 C<sup>δ1</sup>, Tyr159 C<sup>α</sup> | Van der Waals |
| Thr30 O<sup>γ1</sup> | Thr163 O<sup>γ1</sup> | Hydrogen bond |
| Thr30 C<sup>γ2</sup> | Ala158 C<sup>β</sup> | Van der Waals |
| Tyr32 C<sup>ε1</sup> | Gln155 O | Van der Waals |
| Tyr32 C<sup>ζ</sup> | Gln155 O | Van der Waals |
| Tyr32 O<sup>η</sup> | Gln155 C | Van der Waals |
| Tyr32 O<sup>η</sup> | Gln155 O | Hydrogen bond |
| Tyr51 O | Ala158 C<sup>β</sup> | Van der Waals |
| Tyr51 C<sup>γ</sup> | Gln155 C<sup>γ</sup> | Van der Waals |
| Tyr51 C<sup>δ1</sup> | Gln155 N, Gln155 C<sup>γ</sup> | Van der Waals |
| Tyr51 C<sup>ε1</sup> | Glu154 C<sup>β</sup> | Van der Waals |
| Tyr51 O<sup>η</sup> | His151 C<sup>β</sup>, His151 C<sup>γ</sup>, His151 C<sup>δ2</sup> | Van der Waals |
| Ser52 C<sup>β</sup> | Glu154 O | Van der Waals |
| Ser52 O<sup>γ</sup> | Glu154 O | Van der Waals |
| Lys69 C<sup>ε</sup> | Glu166 O<sup>ε2</sup> | Van der Waals |
| Lys69 N<sup>ζ</sup> | Glu166 C<sup>δ</sup> | Van der Waals |
| Lys69 N<sup>ζ</sup> | Glu166 O<sup>ε1</sup>, Glu166 O<sup>ε2</sup> | Salt bridge |
| Leu96 C<sup>γ</sup> | Ala69 C<sup>β</sup> | Van der Waals |
| Leu96 C<sup>δ1</sup> | Ala69 C<sup>β</sup> | Van der Waals |
| Leu96 C<sup>δ2</sup> | Lys66 O, His70 C<sup>ε1</sup> | Van der Waals |
| Ala97 C<sup>β</sup> | Arg65 C<sup>β</sup>, Arg65 C, Arg65 O | Van der Waals |
| Gln98 C<sup>γ</sup> | Arg65 C<sup>δ</sup>, Arg65 N<sup>ε</sup>, Arg65 C<sup>ζ</sup> | Van der Waals |
| Gln98 C<sup>δ</sup> | Arg65 N<sup>η1</sup>, Arg65 N<sup>ε</sup>, Arg65 C<sup>ζ</sup>, Arg65 N<sup>η2</sup> | Van der Waals |
| Gln98 O<sup>ε1</sup> | Arg65 N<sup>η1</sup>, Arg65 N<sup>ε</sup>, Arg65 C<sup>ζ</sup> | Van der Waals |
| Gln98 O<sup>ε1</sup> | Arg65 N<sup>η2</sup> | Hydrogen bond |
| Gln98 N<sup>ε2</sup> | Arg65 N<sup>η1</sup> | Van der Waals |

**Supplementary Table 10: TCR-B $\beta$ (chain D) with HLA-A2 (chain A) interactions found in 9TML.** All intermolecular contacts are assigned according to the following distance cutoffs: hydrogen bonds $\leq 3.4$ Å, salt bridges $\leq 4.5$ Å, Van der Waals $\leq 4.0$ Å.

| TCR-B $\beta$ | HLA-A2 | Bond type |
| --- | --- | --- |
| Trp32 C<sup>52</sup> | Thr73 O<sup>71</sup> | Van der Waals |
| Trp32 C<sup>53</sup> | Gln72 O<sup>61</sup> | Van der Waals |
| Trp32 C<sup>72</sup> | Thr73 O<sup>71</sup> | Van der Waals |
| Arg51 C<sup>5</sup> | Val76 C<sup>71</sup> | Van der Waals |
| Arg51 N<sup>72</sup> | Val76 C<sup>71</sup> | Van der Waals |
| Ser52 O<sup>7</sup> | Val76 C<sup>72</sup> | Van der Waals |
| Gly99 C<sup>α</sup> | Gln155 O<sup>61</sup> | Van der Waals |
| Gly99 C | Gln155 O<sup>61</sup> | Van der Waals |
| Gly100 N | Gln155 C<sup>7</sup>, Gln155 C<sup>6</sup> | Van der Waals |
| Gly100 N | Gln155 O<sup>61</sup> | Hydrogen bond |
| Gly100 C<sup>α</sup> | Gln155 C<sup>6</sup>, Gln155 O<sup>61</sup> | Van der Waals |
| Gly100 C | Gln155 O<sup>61</sup> | Van der Waals |
| Ala101 N | Gln155 O<sup>61</sup> | Van der Waals |

**Supplementary Table 11: TCR-B $\alpha$ (chain C) with GPC3-9-mer (chain P) interactions found in 9TML.** All intermolecular contacts are assigned according to the following distance cutoffs: hydrogen bonds $\leq 3.4$ Å, salt bridges $\leq 4.5$ Å, Van der Waals $\leq 4.0$ Å.

| TCR-B $\alpha$ | GPC3-9-mer | Bond type |
| --- | --- | --- |
| Tyr27 C<sup>ε2</sup> | Glu4 O<sup>ε2</sup> | Van der Waals |
| Tyr27 C<sup>ζ</sup> | Glu4 O<sup>ε2</sup> | Van der Waals |
| Tyr27 O<sup>η</sup> | Phe1 C<sup>ζ</sup>, Glu4 C<sup>γ</sup>, Glu4 C<sup>δ</sup> | Van der Waals |
| Tyr27 O<sup>η</sup> | Glu4 O<sup>ε2</sup> | Hydrogen bond |
| Gly29 C<sup>α</sup> | Phe1 C<sup>ε1</sup>, Phe1 C<sup>ζ</sup>, Glu4 O<sup>ε2</sup> | Van der Waals |
| Gly29 C | Glu4 O<sup>ε2</sup> | Van der Waals |
| Thr30 N | Glu4 O<sup>ε2</sup> | Hydrogen bond |
| Thr30 N | Glu4 C<sup>δ</sup>, Glu4 O<sup>ε1</sup> | Van der Waals |
| Thr30 C<sup>α</sup> | Glu4 O<sup>ε2</sup> | Van der Waals |
| Thr30 O | Glu4 O<sup>ε2</sup> | Van der Waals |
| Thr30 C<sup>β</sup> | Glu4 O<sup>ε2</sup>, Glu4 C<sup>δ</sup>, Glu4 O<sup>ε1</sup> | Van der Waals |
| Thr30 O<sup>γ1</sup> | Glu4 O<sup>ε2</sup>, Glu4 C<sup>δ</sup> | Van der Waals |
| Thr30 O<sup>γ1</sup> | Glu4 O<sup>ε1</sup> | Hydrogen bond |
| Leu96 C<sup>β</sup> | Glu4 C<sup>α</sup>, Glu4 C, Glu4 O | Van der Waals |
| Leu96 C<sup>δ2</sup> | Ala3 O, Glu4 C<sup>α</sup>, Glu4 C, Leu5 N | Van der Waals |

**Supplementary Table 12: TCR-B $\beta$ (chain D) with GPC3-9-mer (chain P) interactions found in 9TML.** All intermolecular contacts are assigned according to the following distance cutoffs: hydrogen bonds $\leq 3.4$ Å, salt bridges $\leq 4.5$ Å, Van der Waals $\leq 4.0$ Å.

| TCR-B $\beta$ | GPC3-9-mer | Bond type |
| --- | --- | --- |
| Trp32 C$^{\delta 1}$ | Asp8 O$^{\delta 2}$ | Van der Waals |
| Trp32 N$^{\epsilon 1}$ | Asp8 C$^{\gamma}$, Asp8 O$^{\delta 1}$ | Van der Waals |
| Trp32 N$^{\epsilon 1}$ | Asp8 O$^{\delta 2}$ | Hydrogen bond |
| Trp32 C$^{\epsilon 2}$ | Asp8 O$^{\delta 1}$, Asp8 O$^{\delta 2}$ | Van der Waals |
| Trp32 C$^{\epsilon 2}$ | Asp8 C$^{\gamma}$, Asp8 O$^{\delta 1}$, Asp8 O$^{\delta 2}$ | Van der Waals |
| Arg51 N$^{\epsilon}$ | Asp8 C$^{\gamma}$ | Van der Waals |
| Arg51 N$^{\epsilon}$ | Asp8 O$^{\delta 1}$, Asp8 O$^{\delta 2}$ | Salt bridge |
| Arg51 C$^{\zeta}$ | Asp8 O$^{\delta 2}$ | Van der Waals |
| Arg51 N$^{\eta 2}$ | Asp8 C$^{\gamma}$ | Van der Waals |
| Arg51 N$^{\eta 2}$ | Asp8 O$^{\delta 2}$ | Salt bridge |
| Gly96 N | Tyr7 C$^{\delta 2}$, Tyr7 C$^{\epsilon 2}$ | Van der Waals |
| Gly96 C$^{\alpha}$ | Ala6 O, Tyr7 C$^{\delta 2}$, Tyr7 C$^{\epsilon 2}$ | Van der Waals |
| Gly96 C | Ala6 O | Van der Waals |
| Thr97 N | Ala6 O | Hydrogen bond |
| Thr97 O | Leu5 C$^{\alpha}$, Leu5 C, Ala6 C$^{\alpha}$, Ala6 C | Van der Waals |
| Thr97 O | Ala6 N, Ala6 O | Hydrogen bond |
| Thr97 C$^{\gamma 2}$ | Ala6 C$^{\beta}$, Ala6 O | Van der Waals |
| Gly98 C$^{\alpha}$ | Glu4 O | Van der Waals |
| Gly98 C | Glu4 O | Van der Waals |
| Gly99 N | Glu4 O | Hydrogen bond |
| Gly99 N | Leu5 C$^{\alpha}$, Leu5 C$^{\beta}$ | Van der Waals |
| Gly99 C$^{\alpha}$ | Glu4 O, Leu5 C$^{\beta}$ | Van der Waals |
| Ala101 N | Tyr7 O$^{\eta}$ | Van der Waals |
| Ala101 C$^{\alpha}$ | Tyr7 O$^{\eta}$ | Van der Waals |
| Ala101 C$^{\beta}$ | Tyr7 O$^{\eta}$ | Van der Waals |

**Supplementary Table 13: HLA-A2 (chain A) with GPC3-9-mer (chain P) interactions found in 9TML.** All intermolecular contacts are assigned according to the following distance cutoffs: hydrogen bonds $\leq 3.4$ Å, salt bridges $\leq 4.5$ Å, Van der Waals $\leq 4.0$ Å.

| HLA-A2 | GPC3-9-mer | Bond type |
| --- | --- | --- |
| Tyr7 Cγ | Leu2 C$\delta^2$ | Van der Waals |
| Tyr7 C$\delta^1$ | Leu2 C$\delta^2$ | Van der Waals |
| Tyr7 C$\epsilon^1$ | Leu2 Cγ, Leu2 C$\delta^2$ | Van der Waals |
| Tyr7 C$\epsilon^2$ | Phe1 N, Phe1 C, Phe1 O | Van der Waals |
| Tyr7 Cζ | Leu2 C$\delta^2$, Phe1 N | Van der Waals |
| Tyr7 Oη | Phe1 Cα, Leu2 N, Phe1 C | Van der Waals |
| Tyr7 Oη | Phe1 N | Hydrogen bond |
| Phe9 C$\epsilon^2$ | Leu2 C$\delta^2$ | Van der Waals |
| Phe9 Cζ | Leu2 C$\delta^2$ | Van der Waals |
| Met45 Cε | Leu2 C$\delta^1$ | Van der Waals |
| Tyr59 Oη | Phe1 N | Van der Waals |
| Glu63 O | Leu2 C$\delta^1$ | Van der Waals |
| Glu63 Cδ | Leu2 N | Van der Waals |
| Glu63 O$\epsilon^1$ | Phe1 Cα, Leu2 Cα, Leu2 Cβ, Leu2 Cγ, Leu2 C$\delta^1$, Phe1 C | Van der Waals |
| Glu63 O$\epsilon^1$ | Leu2 N | Hydrogen bond |
| Glu63 O$\epsilon^2$ | Phe1 Cα, Phe1 Cγ, Phe1 C$\delta^2$, Phe1 C$\epsilon^2$, Leu2 N | Van der Waals |
| Lys66 Cδ | Leu2 O, Ala3 O, Glu4 Cγ | Van der Waals |
| Lys66 Cε | Leu2 Cβ, Leu2 O | Van der Waals |
| Lys66 Nζ | Phe1 Cγ, Phe1 C$\delta^2$, Phe1 C$\delta^1$, Phe1 C$\epsilon^1$, Phe1 C$\epsilon^2$, Phe1 Cζ, Leu2 N, Leu2 Cα, Leu2 C | Van der Waals |
| Lys66 Nζ | Leu2 O | Hydrogen bond |
| Val67 N | Leu2 C$\delta^1$ | Van der Waals |
| Val67 Cα | Leu2 C$\delta^1$ | Van der Waals |
| Val67 Cβ | Leu2 C$\delta^1$ | Van der Waals |
| His70 C$\epsilon^1$ | Ala3 O | Van der Waals |
| Thr73 O$\gamma^1$ | Ala6 Cβ | Van der Waals |
| Thr73 C$\gamma^2$ | Tyr7 C, Asp8 O$\delta^1$, Tyr7 O | Van der Waals |
| Asp77 Cβ | Leu9 C$\delta^1$ | Van der Waals |
| Asp77 Cγ | Leu9 Cγ, Leu9 C$\delta^1$, Leu9 N | Van der Waals |
| Asp77 O$\delta^1$ | Asp8 Cα, Asp8 Cβ, Asp8 C | Van der Waals |
| Asp77 O$\delta^1$ | Leu9 N | Hydrogen bond |
| Asp77 O$\delta^2$ | Leu9 Cγ, Leu9 C$\delta^1$ | Van der Waals |
| Thr80 C$\gamma^2$ | Leu9 O | Van der Waals |
| Leu81 Cγ | Leu9 C$\delta^1$ | Van der Waals |
| Leu81 C$\delta^1$ | Leu9 C$\delta^1$ | Van der Waals |
| Leu81 C$\delta^2$ | Leu9 Cβ, Leu9 C$\delta^1$ | Van der Waals |
| Tyr84 C$\epsilon^2$ | Leu9 O | Van der Waals |
| Tyr84 Cζ | Leu9 O | Van der Waals |
| Tyr84 Oη | Leu9 C, Leu9 O | Van der Waals |
| Tyr84 Oη | Leu9 O | Hydrogen bond |
| Tyr99 Cζ | Ala3 N, Ala3 Cβ | Van der Waals |
| Tyr99 Oη | Leu2 Cα, Leu2 Cβ, Leu2 C$\delta^2$, Leu2 C, Ala3 Cα, Ala3 Cβ | Van der Waals |
| Tyr99 Oη | Ala3 N | Hydrogen bond |
| Tyr116 C$\delta^2$ | Leu9 C$\delta^1$, Leu9 C$\delta^2$ | Van der Waals |
| Tyr123 C$\epsilon^2$ | Leu9 C$\delta^2$ | Van der Waals |
| Thr143 Cα | Leu9 O | Van der Waals |
| Thr143 Cβ | Leu9 O | Van der Waals |
| Thr143 O$\gamma^1$ | Leu9 Cβ, Leu9 Cα, Leu9 C | Van der Waals |
| Thr143 O$\gamma^1$ | Leu9 O | Hydrogen bond |
| Thr143 C$\gamma^2$ | Leu9 C$\delta^2$, Leu9 Cα, Leu9 O | Van der Waals |
| Lys146 Nζ | Leu9 C, Leu9 O | Van der Waals |
| Lys146 N<sup>ε</sup> | Leu9 O | Hydrogen bond |
| Trp147 C<sup>δ1</sup> | Asp8 O | Van der Waals |
| Trp147 N<sup>ε1</sup> | Tyr7 C<sup>β</sup>, Asp8 C | Van der Waals |
| Trp147 N<sup>ε1</sup> | Asp8 O | Hydrogen bond |
| Trp147 C<sup>ε2</sup> | Asp8 O | Van der Waals |
| Trp147 C<sup>ε2</sup> | Tyr7 O, Asp8 O, Leu9 C<sup>δ2</sup> | Van der Waals |
| Trp147 C<sup>η2</sup> | Leu9 C<sup>δ2</sup> | Van der Waals |
| Ala150 C<sup>β</sup> | Tyr7 C<sup>ε2</sup> | Van der Waals |
| Val152 C<sup>γ2</sup> | Tyr7 C<sup>δ1</sup>, Tyr7 C<sup>γ</sup> | Van der Waals |
| Gln155 C<sup>β</sup> | Leu5 C<sup>δ1</sup> | Van der Waals |
| Gln155 C<sup>δ</sup> | Tyr7 C<sup>ε1</sup>, Tyr7 O<sup>η</sup> | Van der Waals |
| Gln155 O<sup>ε1</sup> | Tyr7 C<sup>ε1</sup>, Tyr7 C<sup>ε</sup> | Van der Waals |
| Gln155 O<sup>ε1</sup> | Tyr7 O<sup>η</sup> | Hydrogen bond |
| Gln155 N<sup>ε2</sup> | Tyr7 C<sup>ε1</sup>, Tyr7 C<sup>ε</sup> | Van der Waals |
| Gln155 N<sup>ε2</sup> | Tyr7 O<sup>η</sup> | Hydrogen bond |
| Leu156 C<sup>δ2</sup> | Leu5 C<sup>δ1</sup>, Leu5 C<sup>δ2</sup> | Van der Waals |
| Tyr159 C<sup>δ1</sup> | Glu4 O<sup>ε1</sup>, Ala3 C<sup>α</sup> | Van der Waals |
| Tyr159 C<sup>ε1</sup> | Leu2 C, Leu2 O, Phe1 O, Ala3 N, Ala3 C<sup>α</sup> | Van der Waals |
| Tyr159 C<sup>ε</sup> | Phe1 O, Ala3 N, Ala3 C<sup>α</sup> | Van der Waals |
| Tyr159 O<sup>η</sup> | Leu2 C, Phe1 C, Ala3 N | Van der Waals |
| Tyr159 O<sup>η</sup> | Phe1 O | Hydrogen bond |
| Thr163 C<sup>γ2</sup> | Phe1 C<sup>δ1</sup>, Phe1 C<sup>ε1</sup> | Van der Waals |
| Trp167 C<sup>β</sup> | Phe1 C<sup>β</sup> | Van der Waals |
| Trp167 C<sup>γ</sup> | Phe1 C<sup>β</sup>, Phe1 C<sup>γ</sup> | Van der Waals |
| Trp167 C<sup>δ1</sup> | Phe1 C<sup>β</sup>, Phe1 C<sup>γ</sup> | Van der Waals |
| Trp167 C<sup>δ2</sup> | Phe1 C<sup>β</sup>, Phe1 C<sup>γ</sup>, Phe1 C<sup>δ2</sup> | Van der Waals |
| Trp167 N<sup>ε1</sup> | Phe1 C<sup>γ</sup>, Phe1 C<sup>δ2</sup>, Phe1 C<sup>δ1</sup> | Van der Waals |
| Trp167 C<sup>ε2</sup> | Phe1 C<sup>β</sup>, Phe1 C<sup>γ</sup>, Phe1 C<sup>δ2</sup>, Phe1 C<sup>ε2</sup> | Van der Waals |
| Trp167 C<sup>ε2</sup> | Phe1 C<sup>δ2</sup>, Phe1 C<sup>ε2</sup> | Van der Waals |
| Trp167 C<sup>η2</sup> | Phe1 C<sup>δ2</sup> | Van der Waals |
| Tyr171 C<sup>ε2</sup> | Phe1 N | Van der Waals |
| Tyr171 C<sup>ε</sup> | Phe1 N | Van der Waals |
| Tyr171 O<sup>η</sup> | Phe1 C<sup>α</sup>, Phe1 C<sup>β</sup> | Van der Waals |
| Tyr171 O<sup>η</sup> | Phe1 N | Hydrogen bond |

**Supplementary Table 14. Kinetic constants for TCR-A and TCR-B binding to HLA-A2:GPC3-9-mer.**

Binding kinetic constants for TCR-A and TCR-B.

| TCR | $K_D$ ($\mu\text{M}$) | $k_{on}$ ($\text{M}^{-1} \text{s}^{-1}$) | $k_{off}$ ($\text{s}^{-1}$) |
| --- | --- | --- | --- |
| TCR-A | $382 \pm 79$ | $1265 \pm 140$ | $0.477 \pm 0.047$ |
| TCR-B | $312 \pm 12$ | $761 \pm 224$ | $0.236 \pm 0.061$ |

*Abbreviations:* $K_D$, equilibrium dissociation constant; $k_{on}$, association rate constant; $k_{off}$, dissociation rate constant; Values are presented as mean ± standard deviation.

**Supplementary Table 15. Conjugate residence times by kinetic profile.** Statistical comparison of conjugate formation percentages between target (HepG2) and control (EO771) tumor cell lines for TCR-A (**a**) and TCR-B (**b**) T cells over time. Statistical significance was determined using a two-way ANOVA. ns, not significant; \*p < 0.05; \*\*p < 0.01.

| **TCR-A/HepG2 vs**<br>**TCR-A/EO771** | **0 min** | **10 min** | **30 min** | **60 min** |
| --- | --- | --- | --- | --- |
| P value | 0,9851 | 0,0076 | 0,0315 | 0,0307 |
| Significance | ns | ** | * | * |

| **TCR-B/HepG2 vs**<br>**TCR-B/EO771** | **0 min** | **10 min** | **30 min** | **60 min** |
| --- | --- | --- | --- | --- |
| P value | 0,9606 | 0,9918 | 0,0555 | 0,0057 |
| Significance | ns | ns | ns | ** |
